# Multiple embryonic sources converge to form the pectoral girdle skeleton in zebrafish

**DOI:** 10.1101/2023.07.14.548949

**Authors:** Shunya Kuroda, Robert L. Lalonde, Thomas A. Mansour, Christian Mosimann, Tetsuya Nakamura

**Author notes:** Present address: Institute for Frontier Science Initiative, Kanazawa University, Kakuma-machi, Kanazawa 920-1164, Japan (Shunya Kuroda). Correspondence to: Shunya Kuroda; ORCID: 0000-0003-4820-2165; and Tetsuya Nakamura; ORCID: 0000-0003-0183-1685.

## Abstract

The morphological transformation of the pectoral/shoulder girdle is fundamental to the water-to-land transition in vertebrate evolution. Although previous studies have resolved the embryonic origins of the tetrapod shoulder girdle, those of the fish pectoral girdle remain uncharacterized, creating a gap in the understanding of girdle transformation mechanisms from fish to modern tetrapods. Here, we identified the embryonic origins of the pectoral girdle of zebrafish (*Danio rerio*), including the cleithrum as an ancestral pectoral girdle element lost in extant tetrapods. Our combinatorial approach of photoconversion and genetic cell lineage tracing mapped that cleithrum development combines three adjoining embryonic populations: cranial neural crest cells and lateral plate mesoderm-derivatives (trunk lateral plate mesoderm and cardiopharyngeal mesoderm-associated cells). The topographical position of the cleithrum at the head/trunk interface is a shared characteristic among cleithrum-bearing fish, thus its multiple embryonic origins are likely a conserved feature. Moreover, a comparison of the pectoral girdle progenitors between aquatic fish and extant amniotes suggests that cleithrum loss is associated with the disappearance of its unique developmental environment by the insertion of the neck lateral plate mesoderm into the head/trunk interface. Overall, our study establishes an embryological framework for pectoral/shoulder girdle formation and their evolutionary trajectories from their origin in water to diversification on land.

## Introduction

The pectoral girdle anchors the pectoral appendage (pectoral fin or forelimb) to the body wall in jawed vertebrates (Goodrich, 1930). The pectoral girdle comprises of dermal bones and cartilage or their replacement bones (endoskeleton), while cartilaginous fishes secondarily lost the dermal components (McGonnell, 2001). In bony vertebrates with paired fins living in water (sarcopterygians and actinopterygians), the cleithrum (Gegenbaur, 1895), one of the pectoral dermal bones, is both functionally and anatomically a major component. As vertebrates transitioned onto land, the relative size of the cleithrum in the pectoral girdle began to decrease and eventually disappeared independently in multiple lineages leading to extant tetrapods (Gess & Ahlberg, 2018; Goodrich, 1930; Matsuoka et al., 2005). Consequently, except for some anuran amphibians (Jollie, 1962; Romer & Parsons, 1986) and possibly turtles (Lyson et al., 2013), no extant tetrapods form the cleithrum, and enlarged endoskeletal elements are instead the dominant component of the pectoral girdle in tetrapods (Romer, 1997). In contrast to these accumulating anatomical descriptions, the embryonic changes responsible for the evolutionary loss of the cleithrum and expansion of the endoskeleton remain elusive.

The major embryonic sources of the shoulder girdle in tetrapods have been identified by work in several model systems. Homotopic transplantations between chicken and quail embryos (Nagashima et al., 2016) and genetic cell lineage tracing using *Prx1:Cre* in mice (Heude et al., 2018) revealed that the clavicle, a dermal component of the shoulder girdle, derives mainly from the trunk lateral plate mesoderm (LPM). The same strategy of tissue transplantation and genetic cell lineage tracing using *Mef2c-AHF:Cre* mice (cardiopharyngeal marker) identified that the cardiopharyngeal mesoderm (CPM) also contributes to the medial extremity of the clavicle (Adachi et al., 2020; Nagashima et al., 2016). The CPM or cardiopharyngeal field defines a deeply conserved progenitor field that at least in part within the anterior LPM (Lescroart et al., 2022; Nomaru et al., 2021; Prummel et al., 2020; Wang et al., 2019). Moreover, previous studies discovered that the scapula, an endoskeletal component of the girdle, in axolotls, chickens, and mice derives mainly from the trunk LPM and its dorsal portion from the somites, potentially due to the scapula’s developmental position across these mesodermal boundaries (Durland et al., 2008; Piekarski & Olsson, 2011; Shearman et al., 2011; Valasek et al., 2010). Consequently, the trunk LPM is an important contributor to both dermal and endoskeletal components in tetrapod shoulder girdles, while the CPM and somites contribute to their medial and dorsal margins to some extent.

Neural crest cells are migratory ectodermal cells with broad lineage potential that among its diverse descendant lineages contribute to canonical ectodermal tissues (neurons and gilia) and cranial mesenchymal tissues (skeletons and connective tissues) (reviewed by Etchevers et al., 2019; Fabian & Crump, 2023). Previous work in axolotl tested the potential contribution of neural crest cells to the shoulder girdle, but did not detect any neural crest-derived cells in the shoulder girdle skeletons (Epperlein et al., 2012). In contrast, tissue transplantations in chicken embryos (McGonnell et al., 2001) and genetic lineage tracing using *Sox10:Cre* mice (with *Sox10* being expressed in migratory neural crest cell; Matsuoka et al., 2005) identified that neural crest cells give rise to clavicles. More recent genetic tracing of *Wnt1:Cre* that marks premigratory neural crest cells in mice, however, found no neural crest cell-lineage in the shoulder girdle skeletons (Adachi et al., 2020; Heude et al., 2018). This inconsistency among lineage labeling studies may partially be due to the deployment of isolated *cis*-regulatory elements of different neural crest genes. The distribution of genetically labeled neural crest cell lineages in mice arguably varies depending on the cis-regulatory elements that drive Cre expression (Aoto et al., 2015); consequently, the contribution of the neural crest cells to the shoulder skeletons must be resolved by additional cell lineage tracking such as region-specific photoconversions or tissue transplantations (see an example in Masselink et al., 2016). Nonetheless, in sum, the contribution of neural crest cells to the tetrapod shoulder girdle has not been unambiguously determined and, if any, appears to be minor (Matsuoka et al., 2005; McGonnell et al., 2001).

Revealing the developmental processes forming the pectoral girdle in actinopterygians would provide mechanistic insights into how such intricate tetrapod shoulder development was established. However, in contrast to the accumulated, yet incomplete knowledge of the embryonic origins of tetrapod shoulder girdles, less is known about the developmental lineages contributing to the pectoral girdles of fishes with paired fins. Previous work in zebrafish indicated that the embryonic origin of the dermal pectoral components, including the cleithrum, is the neural crest (Matsuoka et al., 2005). In contrast, *Sox10:Cre*-labeled lineages did not contribute to dermal components of the pectoral girdle in zebrafish (Kague et al., 2012). However, this could be partially due to the deployment of *Sox10* regulatory elements from mice in the used transgenics that might not fully recapitulate zebrafish *sox10* expression or lineage labeling. The endoskeletal scapulocoracoid, a cartilaginous precursor for the scapula and coracoid, seems to be formed within the fin bud mesenchyme in the larval zebrafish (reviewed in Mercader, 2007), yet the embryonic origins of the endoskeletal component have not been fully identified either. Accordingly, the embryonic origins of the dermal and endoskeletal pectoral girdle components in bony fishes remain to be determined.

In zebrafish, the cleithrum lies between the head and trunk structures in the absence of a forming neck in fishes (Figure 1a); the cleithrum posteriorly delineates the most caudal pharyngeal arch and the pericardium, the mesothelium-lined body cavity encapsulating the heart (Figure 1b, c), and anteriorly delineates the trunk lateral body wall where the pectoral fin attaches (Figure 1b, c). Moreover, the cleithrum develops closely associated with the pectoral musculature: cleithrohyoid (see Materials and Methods for nomenclature summary), pectoral fin adductor/abductor, and posterior hypaxial muscles (Figure 1b, c). Embryologically, these anatomical components associated with the cleithrum derive from distinct embryonic sources: the pharyngeal arch mesenchyme from cranial neural crest cells (Kague et al., 2012; Mongera et al., 2013), the pericardium from the LPM including *tbx1*-expressing CPM (Felker et al., 2018; Prummel et al., 2022), the lateral body wall (mesothelium and connective tissues of the posterior hypaxial muscle) and pectoral fin bud from the trunk LPM (Mao et al., 2015; Prummel et al., 2022; Sagarin et al., 2019), and the pectoral musculature from anterior somites (Minchin et al., 2013; Talbot et al., 2019). Thus, given its anatomically unique position, cleithrum development in zebrafish potentially deploys any of these diverse embryonic cell populations (Figure 1a–c).

**Figure 1.**
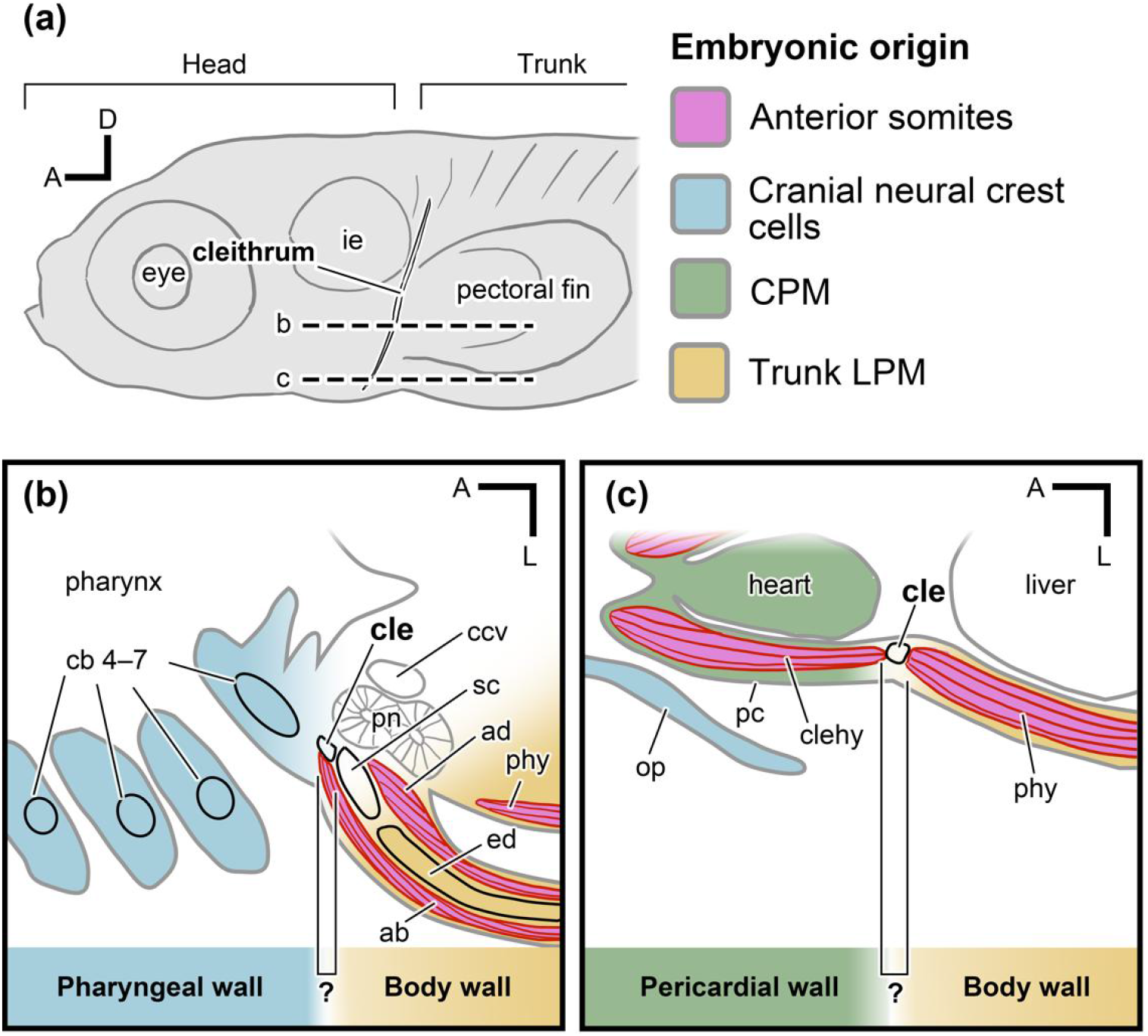
The anatomical position and developmental environment of the pectoral girdle. **(a)** Left lateral view of the larval zebrafish. **(b, c)** Schematic view of horizontal sections obtained from two dorsoventrally different levels as indicated in (a), showing that the larval cleithrum is located at the interface made by multiple embryonic populations: anterior somites (purple), cranial neural crest cells (light blue), CPM (green), and trunk LPM (yellow). The embryonic origins of the pectoral girdle that develops at the boundary of fin-field LPM, cranial neural crest cells, and CPM remains to be identified. Panels (b, c) were drawn after open data sets available on Zebromes (https://neurodata.io/project/zebromes/) originally published in Hildebrand et al., 2017. A, anterior; ab, abductor muscle; ad, adductor muscle; cb 4–7, ceratobranchial 4–7; ccv, common cardinal vein; cle, cleithrum; clehy, cleithrohyoid muscle; CPM, cardiopharyngeal mesoderm; D, dorsal; ed, endoskeletal disc; ie, inner ear; LPM, lateral plate mesoderm; op, operculum; pc, pericardium; phy, posterior hypaxial muscle; pn, pronephros; sc, scapulocoracoid.

Here, by combining region-specific photoconversions and genetic cell lineage analyses, we labeled the cranial neural crest cells, CPM, and trunk LPM to identify the embryonic lineage contributions to the pectoral girdle skeleton in zebrafish. Our results document that the cleithrum develops as a mosaic bone from multiple embryonic populations, while the scapulocoracoid exclusively derives from the fin field-associated LPM. An broad evolutionary comparison of the topographical position of the cleithrum implies that the multiple embryonic origins of the pectoral girdle skeletons are shared features among extinct and extant cleithrum-bearing species. Our data propose that the evolutionary loss of the cleithrum in amniotes might have followed rearrangements of the ancestral developmental environment at the base of cleithrum formation.

## Results

### Anterior somites give rise to pectoral musculature but not to the cleithrum

Previous work found that somites 1–4 are the major contributors to the pectoral musculature in zebrafish (Talbot et al., 2019). Of these, mesenchymal cells derived from somites 1-3 ventrally migrate over the prospective cleithrum area between the pharynx and the pectoral fin bud (Figure 1a-c) and eventually develop into the cleithrohyoid muscle at the bottom of the pharynx (Sagarin et al., 2019; Talbot et al., 2019). We therefore tested whether these somitic mesenchymal cells also contribute to the cleithrum along their migratory route. To this end, we injected Kaede mRNA, encoding a photoconvertible protein, into one-cell stage embryos (Hatta et al., 2006; see Materials and Methods for details) and labeled somites 1–3 by Kaede photoconversion from green to red fluorescence (Kaede-red) at the 10–12 somite stage (mid-segmentation period: Kimmel et al., 1995) (Figures 2a, a’). We then examined the contribution of labeled cells to the pectoral region at 72 hours post-fertilization (hpf) (protruding mouth stage; Kimmel et al., 1995) when the bone matrix of the cleithrum, cartilage, and skeletal muscles become histologically and molecularly evident (see Figure 2—figure supplement 1). Due to laser illumination (405 nm) through the outside of the embryos to target somites, areas of the epidermis and neural tube dorsally and medially adjacent to the target somites, respectively, were also labeled (Figure 2a’). Given the lack of contribution of these non-somitic lineages to skeletogenic mesenchyme (Lee et al., 2013), these non-mesodermal cells were not expected to affect the analysis of somite-derived cell lineages (Figure 2b). Labeled cells derived from the anterior three somites were found in the first to third myotomes (Figure 2c, d) and the cleithrohyoid muscle (Figure 2d). The observed contributions of the anterior three somites to the myotomes and the cleithrohyoid muscle are consistent with previous reports (Minchin et al., 2013; Talbot et al., 2019), supporting the efficacy of our photoconversion. In contrast, no Kaede-red positive cells were found in the cleithrum (n = 0/3; cleithrum positive/all labeled embryos) (Figure 2c, d; Supplementary file 1).

**Figure 2.**
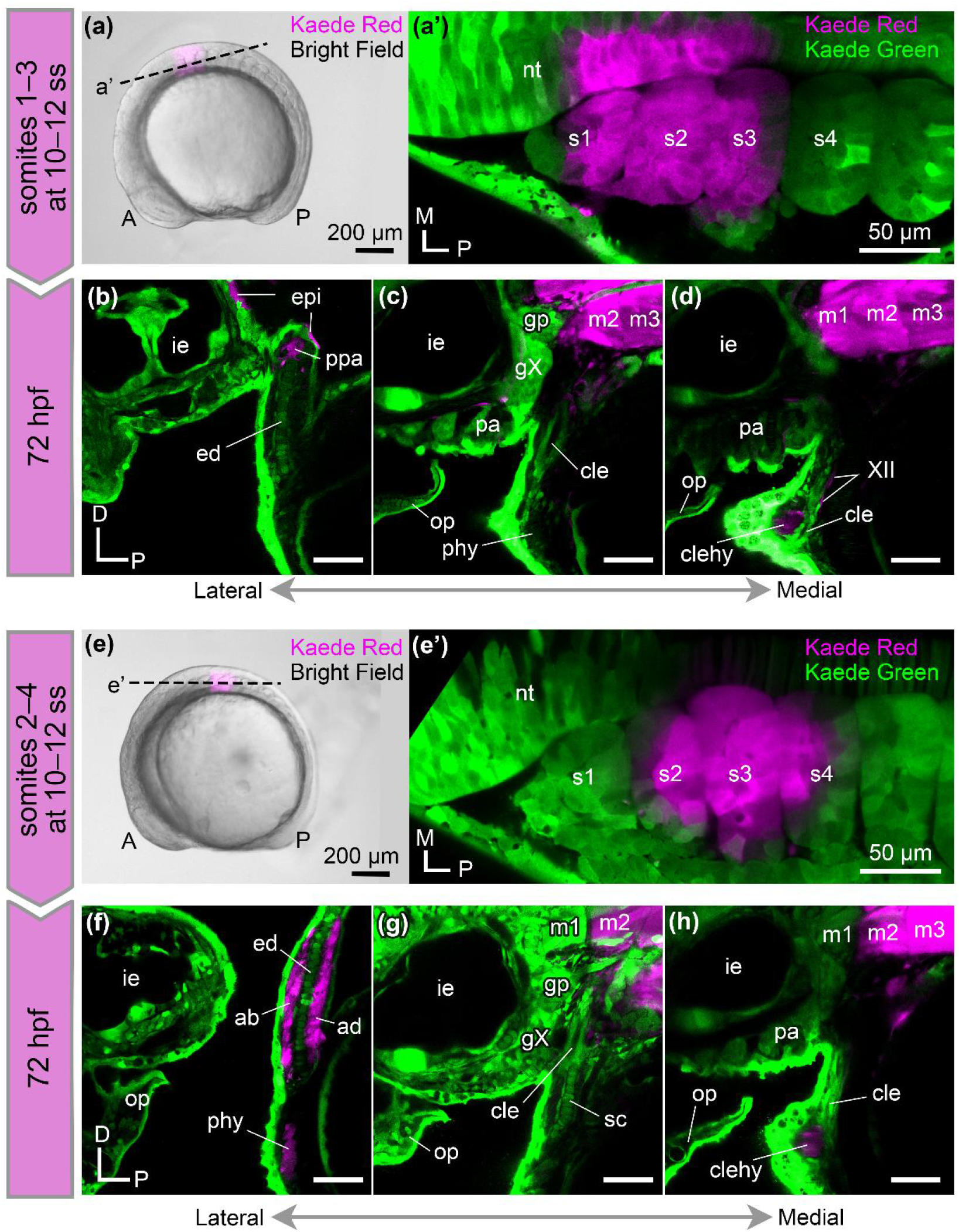
Contribution of anterior somites to pectoral muscles. **(a, a’)** Photoconversion of the anterior three somites and adjacent neural tube at the 10–12 somite stage (ss), viewed from the left lateral side (a) and in horizontal confocal section (a’). Kaede-red is pseudo-colored in magenta. **(b-d)** Confocal parasagittal sections of 72 hpf embryos from lateral– (b) to medial (d) show that labeled cells are in the anterior three myotomes and cleithrohyoid muscle, but not in the cleithrum. **(e, e’)** Photoconversion of the somites 2–4 at the 10–12 ss. **(f-h)** At 72 hpf, parasagittal confocal sections obtained from mediolaterally different levels [lateral (f) to medial (h)] show that labeled cells are in the adductor and abductor fin muscles, posterior hypaxial muscles, myotomes, and cleithrohyoid muscle, but not in the cleithrum. XII, hypoglossal nerve; A, anterior; cle, cleithrum; clehy, cleithrohyoid muscle; ed, endoskeletal disc; epi, epidermis; gp, posterior lateral line ganglion; gX, vagus ganglia; ie, inner ear; op, operculum; P, posterior; pa, pharyngeal arches; phy, posterior hypaxial muscle; ppa, primary pectoral artery; sc, scapulocoracoid; m1–3, myotomes 1–3; nt, neural tube; s1–4, somites 1–4. Scale bars: (a and e) 200 µm, (a’–d, e’–h) 50 µm.

To comprehensively investigate the contribution of the anterior somites, we also photoconverted somites 2–4 at the 10–12 somite stage (Figures 2e, e’). At 72 hpf, labeled cells were found in the pectoral fin adductor/abductor muscles, posterior hypaxial muscles (Figure 2f), myotomes 2–4, and cleithrohyoid muscle (Figure 2 g, h). These distributions also align with previous reports (Minchin et al., 2013; Talbot et al., 2019). Again, no cells labeled with Kaede-red were found in the cleithrum (n = 0/3) or scapulocoracoid (n = 0/3) (Figure 2g, h; Supplementary file 1). In summary, our photoconversion of the anterior somites shows that the first four anterior somites contribute to the pectoral musculature but not to the cleithrum as skeletal element.

### Trunk LPM contributes to the fin mesenchyme, lateral body wall, scapulocoracoid, and cleithrum

Next, to query the contribution of the trunk LPM to the pectoral girdle, we photoconverted the LPM at the prospective pectoral fin region (hereafter “fin-field LPM” as a subpopulation of the trunk LPM) of Kaede-injected embryos at the 10–12 somite stage (Figure 3a, a’). To label the fin-field LPM, we photoconverted the LPM at the level from the first to the third somite (Mao et al., 2015) (Figure 3a, a’). At 72 hpf, the photoconverted cells did not appear in skeletal muscles that arose from the anterior somites (Figure 2), indicating that our photoconversion avoided ectopic labeling of adjacent somites (Figures 3b–d). The labeled cells in the epidermis of the head and pectoral fin likely reflect ectopically labeled epidermal ectoderm (Figure 3b). Labeled cells were also distributed in the endoskeletal disc, the scapulocoracoid (n = 7/7) (Figure 3b; Supplementary file 1), and additionally in the posterior margin of the cleithrum (n = 5/7) (Figure 3c; Supplementary file 1). These data indicate that LPM at the level of the prospective pectoral fin region contributes to the cleithrum.

**Figure 3.**
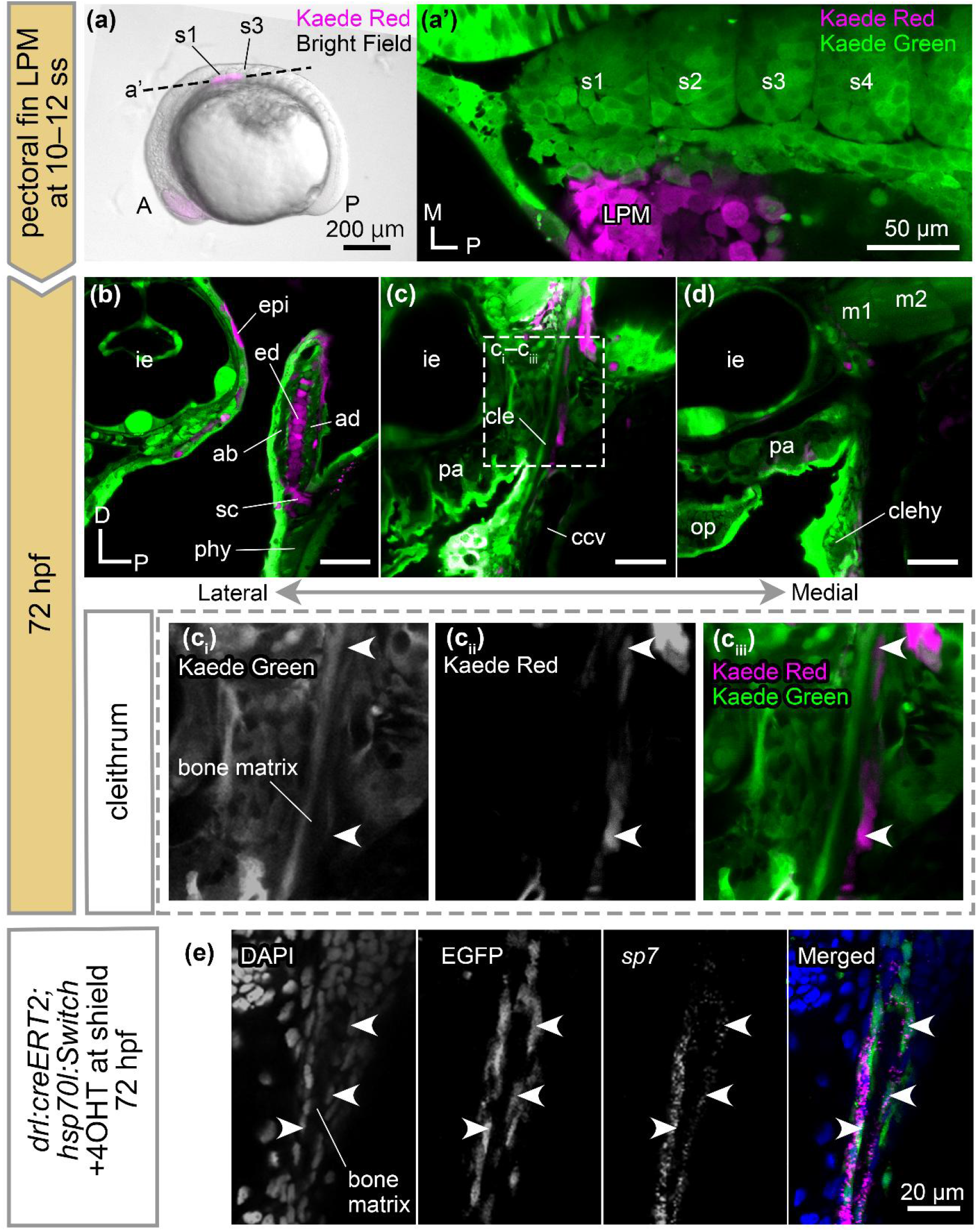
Contributions of the LPM to the pectoral girdle and fin skeleton. **(a, a’)** Photoconversion of the LPM adjacent to somites 1–3 at the 10–12 somite stage (ss), viewed from the left lateral side (a) and in horizontal confocal section (a’). Kaede-red is pseudo-colored in magenta. **(b-d)** At 72 hpf, parasagittal confocal sections obtained from mediolaterally different levels [lateral (b) to medial (d)] show that labeled cells are in the endoskeletal disc, scapulocoracoid, and (ci–ciii) in the posterior half of the cleithrum. (e) Contributions of the genetically labeled *drl:creERT2* lineage cells to the osteoblasts of the cleithrum. Arrowheads in (e) point *drl:creERT2* lineage cells expressing *sp7*. A, anterior; ccv, common cardinal vein; cle, cleithrum; clehy, cleithrohyoid muscle; D, dorsal; ed, endoskeletal disc; epi, epidermis; ie, inner ear; LPM, lateral plate mesoderm; M, medial; m1–2, myotomes 1–2; op, operculum; P, posterior; pa, pharyngeal arches; phy, posterior hypaxial muscle; s1–4, somites 1–4; sc, scapulocoracoid. Scale bars: (a) 200 µm, (a’–d) 50 µm, (e) 20 µm.

To genetically corroborate the contribution of the LPM to the pectoral girdle, we labeled the LPM lineage using *drl:creERT2* transgenic zebrafish, in which the *draculin* (*drl*) regulatory region drives (Z)-4-hydroxytamoxifen (4-OHT)-inducible CreERT2 recombinase expression in the LPM (Mosimann et al., 2015) (see Materials and Methods, and Key Resources Table for detailed strain information). Embryos obtained from crossing *drl:creERT2* and *hsp70l:Switch* reporter (in which EGFP is expressed upon Cre-mediated *loxP* recombination) were treated by 4-OHT from shield stage to 24 hpf and subjected to heat shock at 72 hpf as previously described (Felker et al., 2018; Lalonde et al., 2022) (Figure 3—figure supplement 1). At 72 hpf, EGFP-labeled cells distributed in the cleithrohyoid muscle and the epithelium of the pharyngeal clefts (Figure 3—figure supplement 1a, c, d) are attributable to early-stage *drl* enhancer activity in the somitic lineage and endodermal epithelium of the pharynx, respectively (Mosimann et al., 2015; Sagarin et al., 2019). Consistent with previous studies, EGFP-positive cells were observed in the pectoral fin and aortic arches (Figure 3—figure supplement 1a, b, d) (Mosimann et al., 2015; Lalonde et al., 2022). EGFP-labeled cells were enriched in the cartilage of the endoskeletal disc, scapulocoracoid, and cleithrum (Figure 3-figure supplement 1b, c). Additionally, fluorescent *in situ* hybridization chain reaction (HCR) confirmed that EGFP-positive cells in the cleithrum are osteoblasts expressing an osteoblast marker *sp7* (Figure 3e, Figure 3—figure supplement 1c). Taken together, these results indicate that the region-specific photoconversion and genetic lineage labeling consistently identified the contribution of the LPM to the endoskeletal disc, scapulocoracoid, and cleithrum.

### The LPM containing CPM-associated cells contributes to the cleithrum

To further confirm our finding using *drl:creERT2* based LPM lineage labeling, we sought to refine the spatial photoconversion and genetic labeling into anterior LPM including prospective CPM–derived cells (Felker et al., 2018; Mosimann et al., 2015; Prummel et al., 2022). We photoconverted the ventrolateral head region at the post-otic level (hereafter “ventrolateral head region”), which contains the CPM, at the 10–12 somite stage (Figures 4a). Immediately after photoconversion, we confirmed that the labeled ventrolateral head region contained the prospective CPM, epidermal ectoderm, and ventrolateral half of the post-otic placode (Figure 4a, a’). At 72 hpf, we found labeled cells in the epidermis in the head and pectoral fin, presumably due to ectopic photoconversion of the epidermal ectoderm (Figures 4b-d). The entire vagus ganglia (derivatives of the epibranchial placode: n = 8/8) and the ventral portion of the posterior lateral line ganglion (a derivative of the lateral line placode: n = 7/8) were also labeled (Figure 4c, d; Supplementary file 1), which probably derived from the ectopically labeled ventrolateral half of the post-otic placode at the 10–12 somite stage (Figure 4a’). This result is consistent with the previously reported developmental fate of the post-otic placode (McCarroll et al., 2012; Šestak et al., 2013).

**Figure 4.**
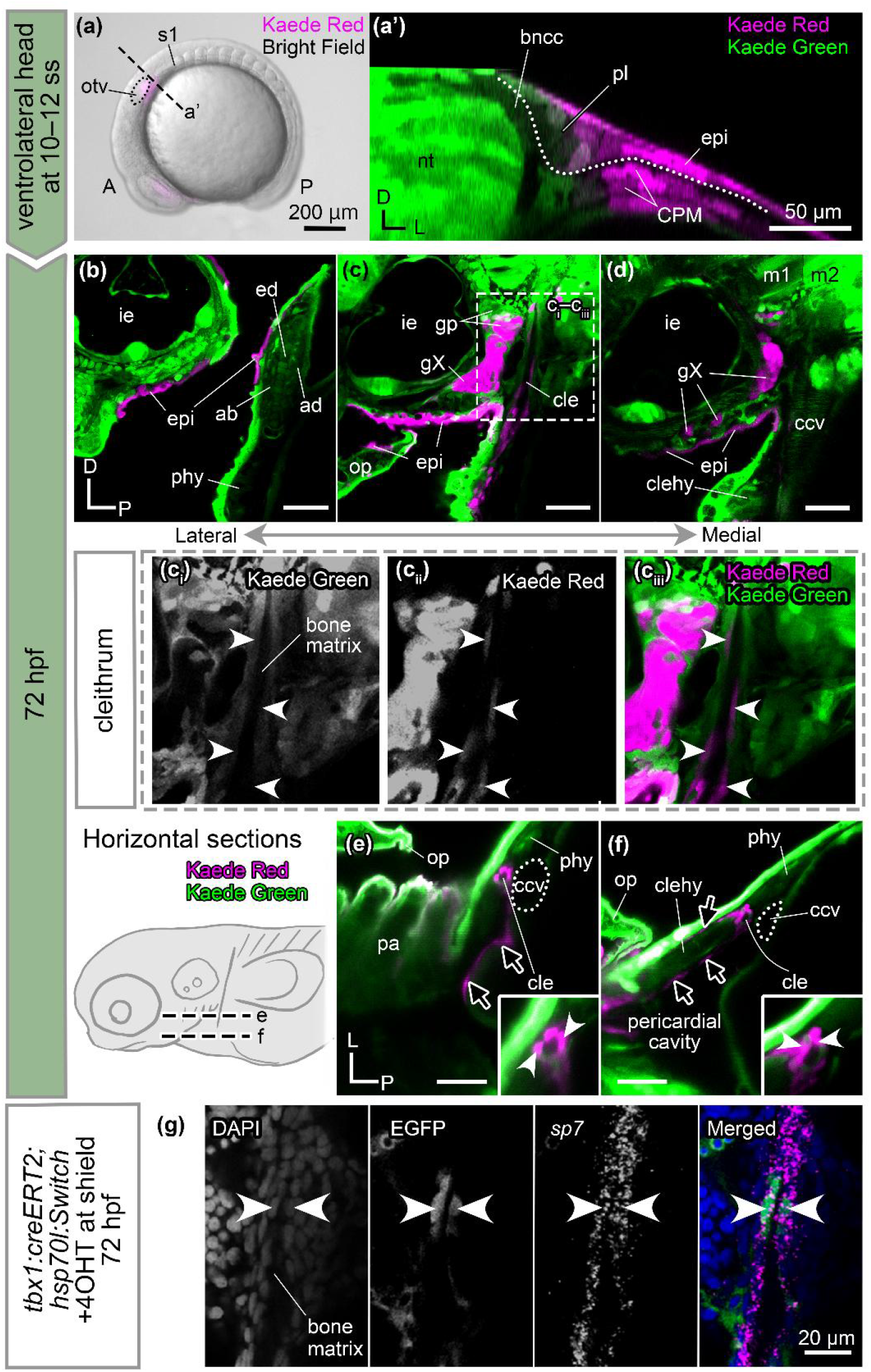
Contributions of CPM to the pectoral girdle and pericardium. **(a, a’)** Photoconversion of the ventrolateral head region at the 10–12 somite stage (ss), viewed from lateral and transverse (a) confocal section (a’). The photoconverted area (pseudo-colored in magenta) in (a’) contains CPM. Dotted line in (a’) depicts the interface between ectodermal epithelium and its underlying tissues. **(b-d)** At 72 hpf, parasagittal confocal sections at mediolaterally different levels [lateral (b) to medial (d)] show that labeled cells are in the ventral part of the posterior lateral line ganglion, vagus nerve ganglia, and (ci–ciii) in the cleithrum. **(e, f)** Horizontal confocal sections at different dorsoventral levels also show labeled cells in the cleithrum (white arrowheads) laterally adjacent to the common cardinal vein (outlined by a dotted line). Labeled cells are also observed in the pericardial wall (black arrows). **(g)** Contribution of the genetically labeled *tbx1:creERT2* lineage cells to osteoblasts of the cleithrum. Arrowheads in (g) point *tbx1:creERT2* lineage cells expressing *sp7*. A, anterior; bncc, branchial neural crest cells; ccv, common cardinal vein; cle, cleithrum; clehy, cleithrohyoid muscle; CPM, cardiopharyngeal mesoderm; D, dorsal; ed, endoskeletal disc; epi, epidermis; gp, posterior lateral line ganglion; gX, vagus nerve ganglia; ie, inner ear; L, lateral; lpm, lateral plate mesoderm; nt, neural tube; op, operculum; otv, otic vesicle; P, posterior; pa, pharyngeal arches; phy, posterior hypaxial muscle; pl, placode; m1–2, myotomes 1–2; s1, first somite. Scale bars: (a) 200 µm, (a’–f) 50 µm, (g) 20 µm.

Labeled cells were also found to delineate the bone matrix of the cleithrum without specific spatial distribution (n = 7/8) (Figure 4c; Supplementary file 1). In horizontal confocal sections, we found that the labeled cells surrounding the bone matrix of the cleithrum (Figure 4e, f; white arrowheads) are contiguous with the mesenchymal cells at the posterior edge of the pericardial wall (Figure 4e, f; black arrows) that itself originates from the LPM including *tbx1*-expressing presumptive CPM (Lalonde et al., 2022; Prummel et al., 2022). Overall, these results indicate that anterior LPM including CPM-associated cells (Figure 4a’) generates osteoblasts contributing to the cleithrum.

To corroborate the result of the anterior LPM and CPM photoconversion, we subsequently performed a genetic cell lineage tracing of *tbx1:creERT2*;*hsp70l:Switch* embryos (see Materials and Methods and Key Resources Table). The T-box transcription factor gene *Tbx1* shows evolutionarily conserved expression patterns in the pharyngeal mesoderm across chordates (Adachi et al., 2018; Onimaru et al., 2011). As such, the zebrafish *tbx1:creERT2* transgene indelibly labels cardiopharyngeal lineages, endoderm in the pharynx, and possibly mandibular neural crest lineages in zebrafish embryos (Felker et al., 2018; Lalonde et al., 2022). We treated embryos obtained from crossing *tbx1:creERT2* and *hsp70l:Switch* zebrafish with 4-OHT from the shield stage to 24 hpf and observed the distribution of EGFP-positive cells at 72 hpf as previously described (Lalonde et al., 2022) (Figure 4—figure supplement 1). Consistent with previous studies (Felker et al., 2018; Lalonde et al., 2022), EGFP-positive cells were observed among mesenchymal cells surrounding the inner ear and in pharyngeal musculature (Figure 4—figure supplement 1a– c). Additionally, labeled cells sparsely delineated the cleithrum bone matrix. HCR *in situ* confirmed that these cells are *sp7*-positive osteoblasts (Figure 4g, Figure 4—figure supplement 1c), demonstrating *tbx1*-positive lineage contribution to the cleithrum. Labeled cells were also found in the epithelium of the pharyngeal clefts (Figure 4—figure supplement 1d), reflecting early *tbx1:creERT2* activity in the endodermal lineage, akin to *Tbx1* in mouse (Arnold et al., 2006; Felker et al., 2018). Taken together, the results of both photoconversion and genetic lineage tracing of the *tbx1:creERT2*-labeled lineage further support a contribution of the anterior LPM and presumptive CPM-associated cells to the zebrafish cleithrum (Figure 4, Figure 4—figure supplement 1).

### Branchial neural crest cells give rise to the pharyngeal mesenchyme and the anterior half of the cleithrum

Several elements of the tetrapod shoulder girdle integrate cells of different lineage origin (Adachi et al., 2020; Durland et al., 2008; Heude et al., 2018; Nagashima et al., 2016; Piekarski & Olsson, 2011; Shearman et al., 2011; Valasek et al., 2010). In our LPM-focused experiments above, we noted highly mosaic lineage labeling in the cleithrum, indicating that the cleithrum might also integrate cells from additional lineage origins. The neural crest contributes to numerous features in the craniofacial skeleton as major evolutionary innovation in vertebrates (Gans & Northcutt, 1983; Kuratani & Ahlberg, 2018). To date, a possible contribution of the neural crest to the cleithrum remains unclear. We therefore used photoconversion to revisit a potential cleithrum contribution of branchial neural crest cells, an evolutionarily conserved, posterior-most cranial neural crest stream in jawed vertebrates (Stundl et al., 2020). We first photoconverted the dorsomedial region of the embryonic head at the post-otic level [hereafter referred to as “dorsomedial head region” corresponding to a previously identified cardiac neural crest population (Sato & Yost, 2003)] in embryos ubiquitously expressing Kaede-green at the 10–12 somite stage (Figures 5a). HCR *in situ* discerned a cell population positive for *foxd3*, a migrating neural crest cell marker (Montero-Balaguer et al., 2006), between the otic vesicle and first somite, confirming that the photoconverted dorsomedial head region includes the branchial neural crest cells (compare Figure 5ai with aii). The adjacent otic vesicle, neural tube, epidermal ectoderm, and dorsomedial portion of the post-otic placode were also ectopically labeled (Figure 5a, ai). The lateral migrating frontier of the branchial neural crest stream at this stage does not reach beyond the placodal region (Figure 5aii) and does not enter into the ventrolateral head region (compare Figure 4a’ with Figure 5aii), indicating that our photoconversions of the branchial neural crest cells (in the dorsomedial head region) and prospective CPM (in the ventrolateral region) were mutually exclusive.

**Figure 5.**
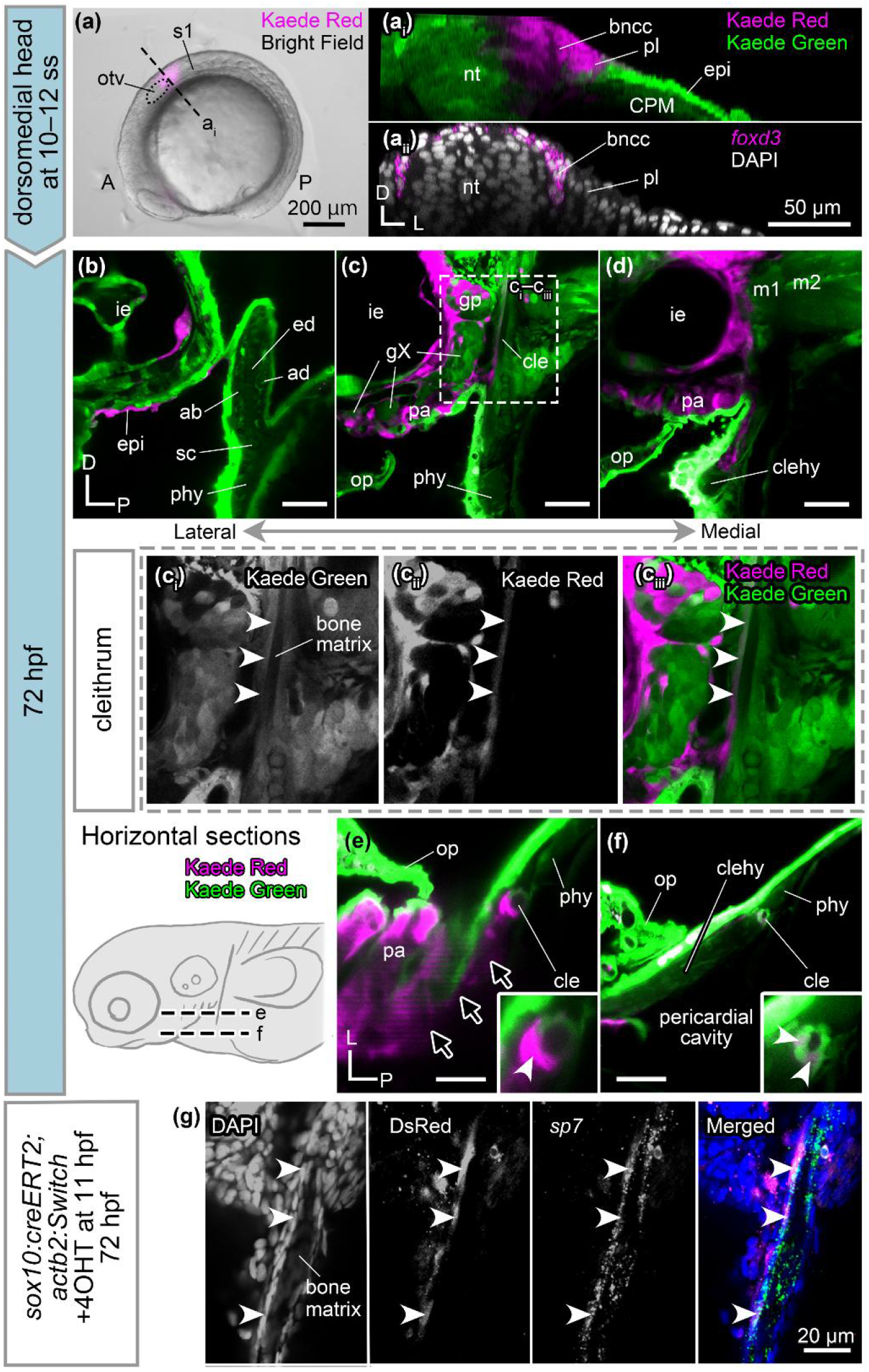
Contributions of cranial neural crest cells to the head and pectoral girdle. **(a, ai)** Photoconversion of the dorsomedial head region at the 10–12 somite stage (ss) labels the branchial neural crest cells and adjacent neural tube, epidermis, and placode. **(aii)** *foxd3 in situ* HCR image obtained from approximately at the same transverse level in (ai). The lateral migratory frontier of branchial neural crest cells does not laterally exceed the level of the placode at this stage. Dotted lines in (ai and aii) depict the interface between ectodermal epithelium and its underlying tissues. **(b-d)** At 72 hpf, parasagittal confocal sections at mediolaterally different levels [lateral (b) to medial (d)] show that labeled cells are in the dorsal part of the posterior lateral line ganglion, mesenchymal cells in the pharyngeal arches, and the anterior half of the cleithrum (ci– ciii). **(e, f)** Horizontal confocal sections at different dorsoventral levels also show labeled cells in the cleithrum (white arrowheads). Labeled cells are continuously distributed from the pharyngeal arch mesenchyme to the cleithrum (black arrows). **(g)** Contribution of the genetically labeled *sox10:creERT2* lineage to osteoblasts of the cleithrum. Arrowheads in (g) point *sox10:creERT2* lineage cells expressing *sp7*. A, anterior; bncc, branchial neural crest cells; cle, cleithrum; clehy, cleithrohyoid muscle; CPM, cardiopharyngeal mesoderm; D, dorsal; ed, endoskeletal disc; epi, epidermis; gp, posterior lateral line ganglion; gX, vagus nerve ganglia; ie, inner ear; L, lateral; lpm, lateral plate mesoderm; nt, neural tube; op, operculum; otv, otic vesicle; P, posterior; pa, pharyngeal arches; phy, posterior hypaxial muscle; pl, placode; m1–2, myotomes 1–2; s1, first somite; sc, scapulocoracoid. Scale bars: (a) 200 µm, (a’–f) 50 µm, (g) 20 µm.

At 72 hpf, Kaede-red labeled cells were found in the posterior wall of the inner ear, epidermis, and posterior lateral line ganglion (Figure 5b–d), consistent with ectopic photoconversion of the otic vesicle, epidermal ectoderm, and dorsolateral half of the post-otic placode, respectively, using our laser-based approach (Figure 5a, ai). Nonetheless, the labeled mesenchymal cells around the vagus ganglia in the posterior pharyngeal arches match previously observed descendants of cranial neural crest cells (Abrial et al., 2017), supporting our photoconversion efficacy (Figure 5c–e). We also found that labeled cells delineate the anterior margin of the cleithrum (n = 10/12) (Figure 5c, white arrowheads in Figure 5e, f, Supplementary table 1) and disseminate toward the labeled pharyngeal arch mesenchyme (black arrows in Figure 5e). These observations indicate a branchial neural crest cell contribution to the cleithrum in addition to our previously established LPM contribution.

As complementary approach and to genetically track the cell lineage of neural crest cells, we performed Cre/*lox*-mediated genetic lineage labeling. We first used *crestin:creERT2* with *ubi:Switch* as loxP reporter (GFP to mCherry change upon Cre activity; see Materials and Methods, and Key Resources Table). The *crestin* regulatory elements are selectively active in the early zebrafish neural crest (Kaufman et al., 2016). In embryos treated with 4-OHT at shield stage and imaged at 96 hpf, we consistently observed mCherry-labeled cells in a pharyngeal arch derivative and recovered individual larvae with mCherry-positive cells in the cleithrum (n = 2/6; Figure 5— figure supplement 1). The overall sparse labeling efficiency with *crestin:creERT2* is consistent with previous reports using this line (Kaufman et al., 2016), yet is consistent with our photoconversion-based observation of neural crest contribution to the cleithrum.

To further corroborate our lineage labeling results, we obtained embryos from crossing *sox10:creERT2* and *actb2:Switch* reporter zebrafish (BFP to DsRed change upon Cre activity; see Materials and Methods, and Key Resources Table). The activity of the zebrafish *sox10* promoter deployed here (Carney et al., 2006) has been reported to have two major phases of expression during zebrafish embryogenesis: a first phase from 11–16 hpf in the premigratory and early migratory neural crest cells and the otic epithelium (Carney et al., 2006; Das & Crump, 2012; Mongera et al., 2013), and a second phase at 48 hpf and after in all chondrocytes regardless of germ layer origins (Dutton et al., 2008; Giovannone et al., 2019). To predominantly label migrating neural crest cells, we activated CreERT2 activity by treating the embryos with 4-OHT from 11 to 24 hpf (Figure 5—figure supplement 2). At 72 hpf, DsRed-positive cells were found in mesenchymal cells in the operculum, pharyngeal arch mesenchyme, epithelium of the inner ear, and cleithrum (Figure 5—figure supplement 2). HCR *in situ* confirmed that DsRed-positive cells expressing *sp7* were localized in only the anterior half of the cleithrum (Figure 5g, Figure 5— figure supplement 2c), consistent with our independent, Kaede-based photoconversion result (Figure 5c). Labeled cells were also discerned sparsely in the cartilages of the endoskeletal disc (Figure 5—figure supplement 2a, b), cells which have previously been reported to show promiscuous *sox10* reporter activity (Howard IV et al., 2021). Overall, both photoconversion and genetic lineage analysis of the *sox10:creERT2* lineage is consistent with contribution of cranial neural crest cells to the anterior half of the cleithrum at the posterior margin of the pharyngeal arch mesenchyme (Figure 5c–g, Figure 5—figure supplement 2).

## Discussion

Our photoconversion and genetic-cell lineage tracing experiments here provide evidence that the zebrafish cleithrum develops as a composite bone of LPM-derived cells (the fin-field LPM and CPM-associated cells) and cranial neural crest cells, while the endoskeletal scapulocoracoid derives solely from the fin-field LPM (Figure 6). The conflicting data of neural crest contribution from previous work (Kague et al., 2012) may be attributed to an activity difference between the mouse *Sox10* enhancer and the here deployed zebrafish *sox10* promoter (Carney et al., 2006), observations with which we further support using *crestin*-based genetic lineage labeling and targeted Kaede photoconversion (Figure 5, Figure 5—figure supplement 1 and 2). Intriguingly, photoconverted fin-field LPM cells were found at the posterior margin of the cleithrum (Figures 3 and 6) while branchial neural crest cells were found only at the anterior margin (Figures 5 and 6). In contrast, *tbx1* lineage-labeled and generally CPM-associated LPM cells distributed throughout the cleithrum (Figures 4 and 6). The spatially confined aggregation of the fin-field LPM and branchial neural crest cells in the cleithrum might result from the position of the cleithrum primordium which lies anteriorly to the pectoral fin bud and posteriorly to the pharynx (Heude et al., 2014). Unlike the single embryonic lineage origin of the zebrafish scapulocoracoid, the endoskeletal scapula in tetrapods arises from two distinct cell populations: LPM and somitic/paraxial mesoderm (Piekarski & Olsson, 2011; Shearman et al., 2011; Valasek et al., 2010). Thus, the somitic contribution into the shoulder girdle could be a derived character in tetrapods due to the dorsal expansion of the scapula. Overall, the zebrafish larval cleithrum develops at the head/trunk interface by assembling topographically close cell populations such as the head (CPM and cranial neural crest cells) and trunk (fin-field LPM) mesenchyme (Figure 6, Figure 6—figure supplement 1a). Together, our data propose a scenario where the cleithrum integrates both LPM– and neural crest cell-derived progenitors are evolutionarily significant interface at the head and trunk developmental programs.

**Figure 6.**
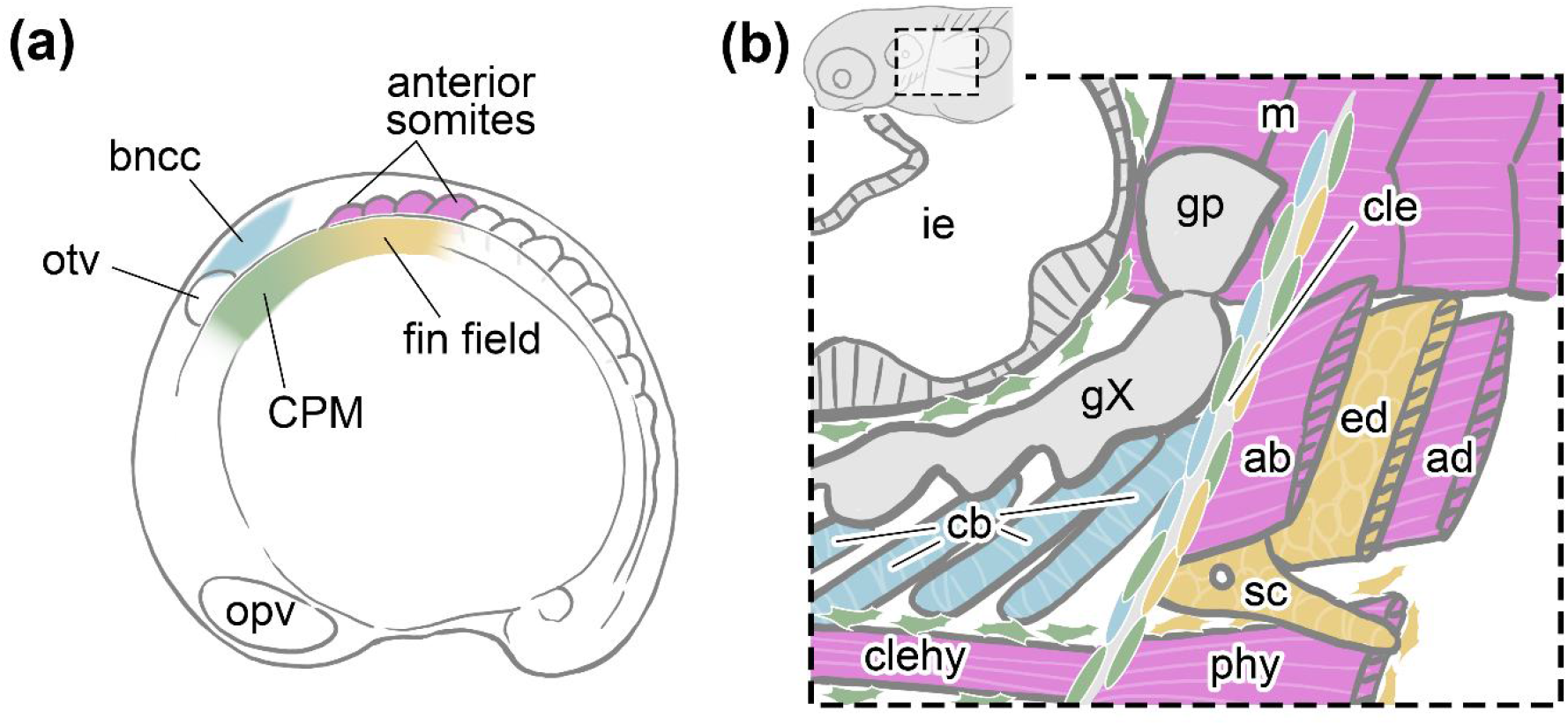
Embryonic origins and environment of the zebrafish pectoral girdle. **(a)** Four embryonic populations reside at the prospective pectoral girdle region at the 10–12 somite stage. They establish the embryonic head/trunk interface in the zebrafish embryo by the pharyngula stage. **(b)** Schematic drawing of the left pectoral region of a zebrafish larva. Based on its embryonic origin, each element follows the same color code as in (a): somites-derived, pink; fin-field LPM-derived, yellow; CPM-derived, green; branchial neural crest cell-derived, light blue. The branchial neural crest cells contribute to ceratobranchial cartilages and anterior ridge of the cleithrum. The CPM gives rise to mesenchymal cells surrounding the cleithrohyoid muscle and is mosaically distributed throughout the cleithrum. The fin-field LPM forms the posterior half of the cleithrum. The scapulocoracoid and endochondral disc originate exclusively from the fin-field LPM. The oval cells surrounding the bone matrix of the cleithrum represent osteoblasts. The distal portion of the pectoral fin elements and the posterior portion of the posterior hypaxial muscle is partially removed (cut surfaces are shaded). ab, abductor muscle; ad, adductor muscle; bncc, branchial neural crest cells; cb, ceratobranchial; cle, cleithrum; CPM, cardiopharyngeal mesoderm; ed, endoskeletal disc; gp, posterior lateral line ganglion; gX, vagus ganglia; ie, inner ear; m, myotomes; opv, optic vesicle; otc, otic vesicle; phy, posterior hypaxial muscle; sc, scapulocoracoid.

The head/trunk interface generates multiple structures, including the neck, that grants independent mobility to the skull from the pectoral girdle (Trinajstic et al., 2013). Neck musculoskeletal components have been reported to originate from a mixture of multiple embryonic cell populations in a variety of jawed vertebrates. In mice, neck musculatures, such as the trapezius, sternocleidomastoid, and infrahyoid muscles, contain connective tissues derived from a combination of cranial neural crest cells and LPM (Adachi et al., 2020; Heude et al., 2018). Moreover, the gill arch skeletons in cartilaginous fishes and amphibians and the laryngeal skeleton in amniotes develop as composites of cranial neural crest cells and CPM (Evans & Noden, 2006; Sefton et al., 2015; Sleight & Gillis, 2020; Tabler et al., 2017). Likewise, the cleithrum develops between the pharyngeal arches and pectoral fin, deploying distinct cellular sources and, possibly, developmental genetic programs. However, the functional contributions or properties of these mixed embryonic origins in neck and shoulder girdle evolution remain unknown.

The detailed analysis of the embryonic origins of the cleithrum and scapulocoracoid illuminates the intricate developmental mechanisms of the pectoral girdle in bony fishes. Several genetic mutant zebrafish with a complete loss of the pectoral fin and scapulocoracoid develop a completely or almost normal cleithrum (reviewed in Mercader, 2007), suggestive of a certain degree of ontogenetic independency between the cleithrum and scapulocoracoid, at least during early pectoral girdle development. We propose that this independence is associated with their divergent embryonic origins that we addressed in this study (Figure 6). Concomitantly, the fin-field LPM, the shared embryonic population between the two types of pectoral skeletons, could be responsible for the integration of developmental processes of these skeletal elements. Indeed, no extant or extinct animals possess the cleithrum detached from the scapula or coracoid, implying interdependent developmental programs. Overall, the investigation of fish pectoral girdle origins explains a wide range of phenotypes that have been difficult to interpret by genetics or genomics alone.

Multiple embryonic origins of the cleithrum provides novel insights into the evolution of the pectoral girdle in fin-bearing gnathostomes. The head/trunk interface where the zebrafish cleithrum develops (Figure 6) is embryologically defined as the distribution boundary between the head and trunk mesenchyme (Kuratani, 1997). Importantly, anatomical structures around the interface (e.g., circumpharyngeal ridge, common cardinal vein, hypoglossal nerve, and hypoglossal cord) are highly preserved across all jawed vertebrates (Higashiyama et al., 2016; Hirasawa & Kuratani, 2013; Kuratani, 1997; Naumann et al., 2017); thus, the relative position of cleithrum with these anatomical structures serves as a means to infer the embryonic environment of the cleithrum in extinct and extant species. For instance, in Australian lungfish (*Neoceratodus*), an extant fin-bearing sarcopterygian, the topographical relationship among the circumpharyngeal ridge, common cardinal vein, pronephros, and cleithrum is comparable to that of zebrafish (see Greil, 1913; Hirasawa et al., 2021 and compare with Figure 1b). This conserved topographical relationship was likely established and fixed in the course of the stem-group gnathostomes (Adachi et al., 2018; Higashiyama et al., 2016; Trinajstic et al., 2022). Moreover, their tight topographical association even dates back to extinct jawless gnathostomes, osteostracans that possess early paired appendages and dermal pectoral girdle (Janvier et al., 1991; discussed in Adachi et al., 2016). Thus, despite the anatomically derived characteristics of the teleost pectoral girdle among actinopterygians (Jollie, 1962; Kingsley, 1917), the embryonic environment of the cleithrum identified in this study (Figure 6) seems to be conserved across fishes bearing dermal pectoral girdles. We propose that, since its emergence, the pectoral appendage position has been confined to the posterior edge of the head and heart due to developmental constraints on the cleithrum (Janvier, 1996; Trinajstic et al., 2022).

Early Synapsida and Sauropsida, such as *Edaphosaurus* (stem Synapsida; Williston, 1925) and Procolophonoidea (stem Sauropsida; MacDougall et al., 2013), retained the cleithrum even in their entirely terrestrial habitats. However, along with increases in the cervical vertebral number and extension of the neck (Hirasawa & Kuratani, 2013; Müller et al., 2010), their crown-groups, including all extant amniotes, completely lost the cleithrum (Jollie, 1962; Romer, 1997). In extant amniotes, the long neck integrates cells from a specialized trunk LPM region, namely the neck LPM (Lours & Dietrich, 2005), which is located between the head/trunk interface and forelimb bud (Figure 6—figure supplement 1b, c). A previous study in chicken embryos demonstrated that the neck LPM is distinct from other portions of the trunk LPM in its developmental incompetency to respond to environmental signals, such as Fgf8 from the ectoderm (Lours & Dietrich, 2005).

Similarly, in mouse embryos, the neck LPM is identified as the “cervical lateral body wall” (Hirasawa et al., 2016). Its expansion during development is critical for the posterior relocation of the heart and providing the mesenchymal environment for the diaphragm, which is a mammalian specific structure (Figure 6—figure supplement 1c) (Hirasawa et al., 2016; Sefton et al., 2018; Sefton et al., 2022). Although it remains unknown whether birds and mammals share the common evolutionary origin of the neck LPM, the insertion and specialization of the neck LPM into the head/trunk interface separated the head mesenchyme and forelimb LPM, that gives rise to the scapula and coracoid (Capellini et al., 2010; Nagashima et al., 2016). Thus, the evolution of the neck LPM was accompanied by rearrangement of the embryonic head/trunk interface, likely leading to the disappearance of the ancestral developmental environment of the amniote cleithrum (Figure 6—figure supplement 1).

Recent studies have demonstrated that embryonic sources and morphological homology can be evolutionarily decoupled (Fabian & Crump, 2023; Piekarski & Olsson, 2011; Schneider, 1999; Sleight & Gillis, 2020; Teng et al., 2019). If this is the case in pectoral/shoulder girdle evolution, the embryonic sources for the cleithrum could vary in different fish species. Moreover, contrary to the above discussion, the cleithrum may not simply be a lost component in extant tetrapods; its developmental program may be maintained in the shoulder girdle even with its distinct embryonic environment. Further studies of the cellular trajectories and molecular mechanisms underlying these cell differentiations in various cleithrum-bearing fishes are warranted to determine the possibility of decoupling of ontogenetic origins and homological traits over the course of pectoral/shoulder girdle evolution.

## Materials and Methods

### Key resources table

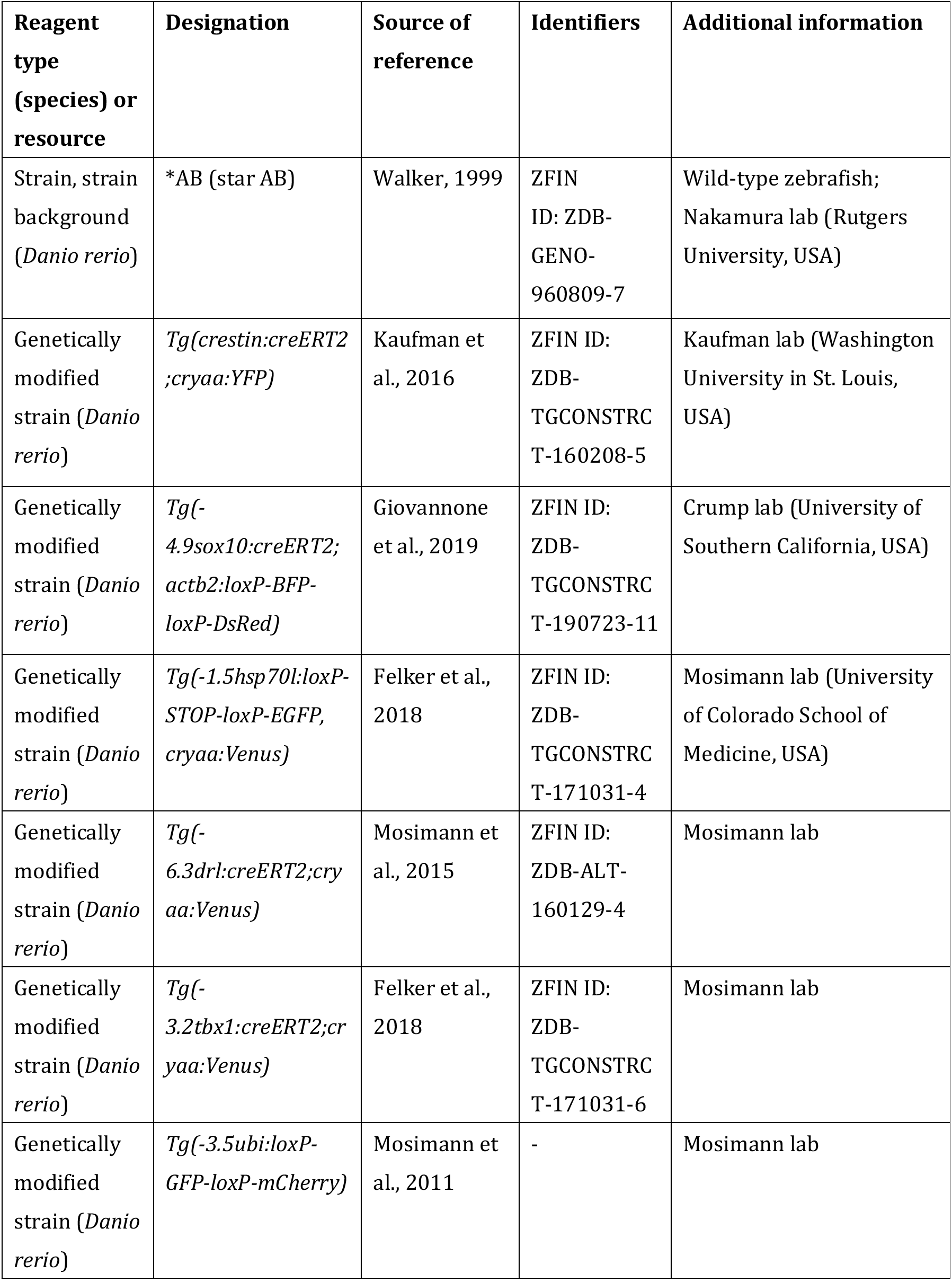
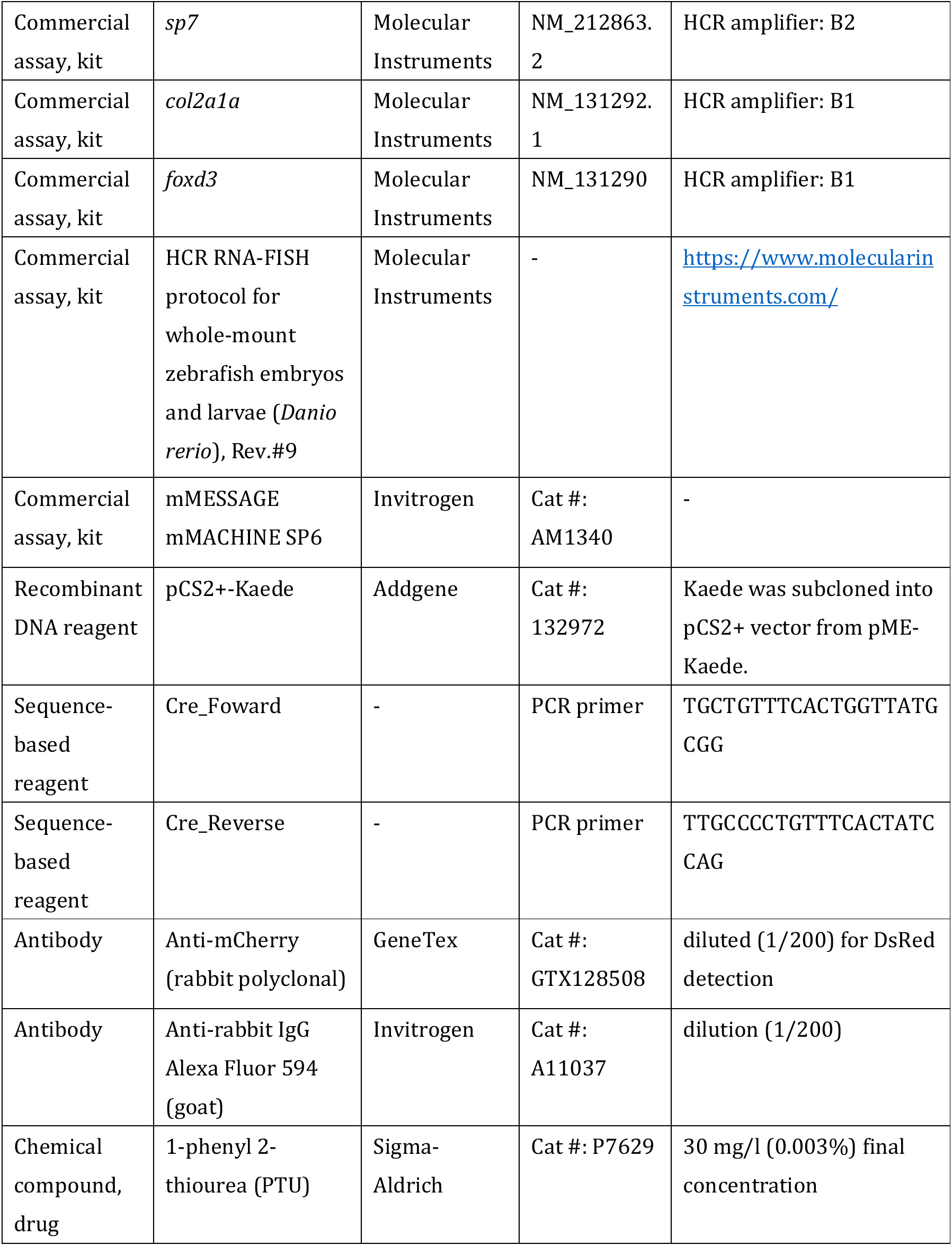
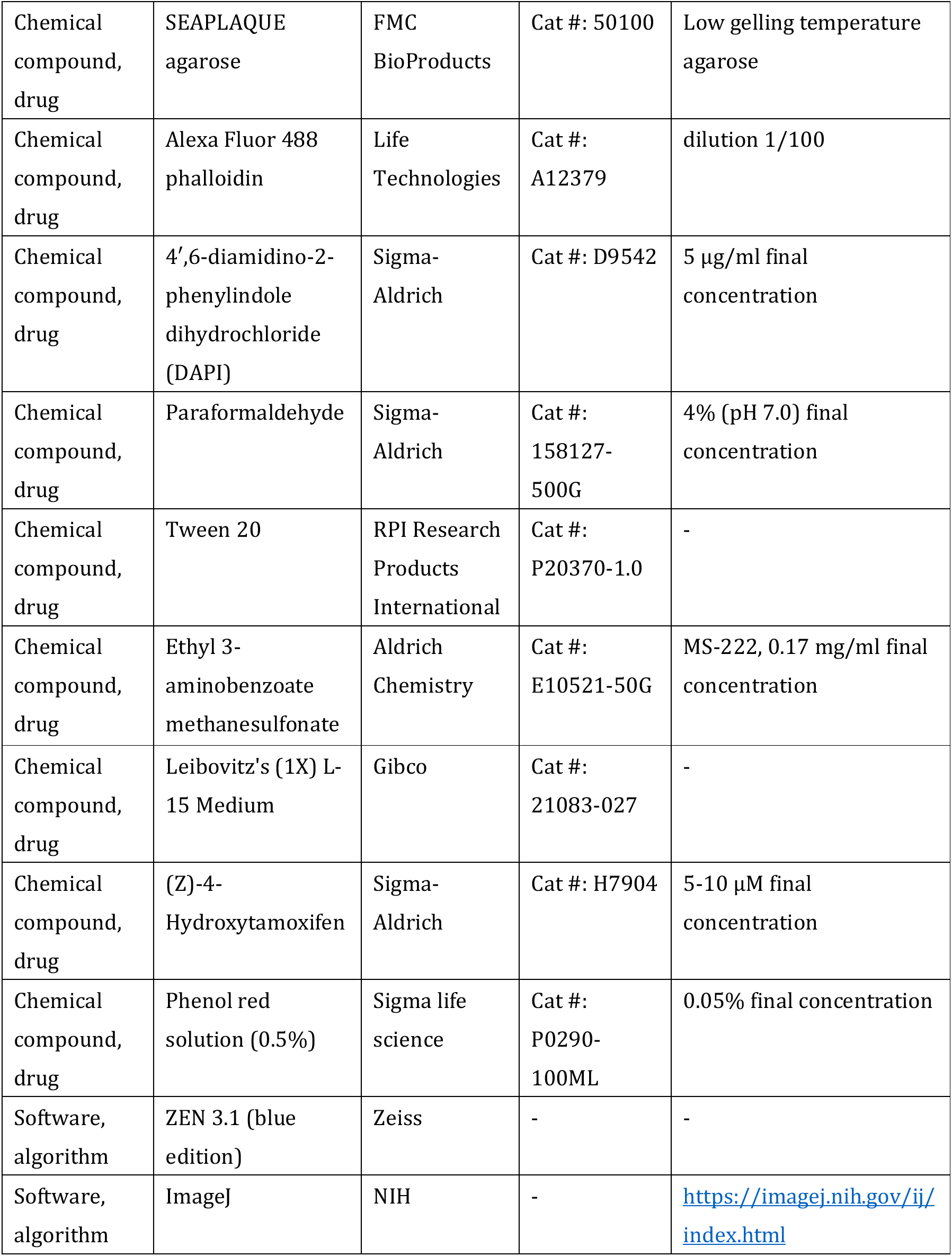
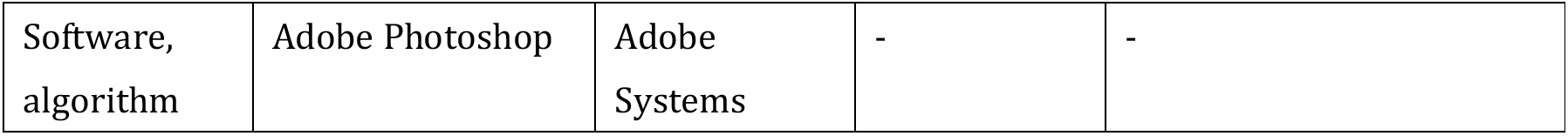

### Zebrafish lines

Animal husbandry and experiments were carried out in accordance with the protocol approved by Institutional Animal Care and Use Committee (IACUC) of Rutgers University (protocol #:201702646) and of the University of Colorado School of Medicine (protocol #00979). Adult zebrafish [*Danio rerio* (Hamilton, 1822)] were kept at 28.5°C on a 14 hours light/10 hours dark cycle. Wild-type zebrafish embryos were obtained from intercross of *AB (star-AB) line. Zebrafish transgenic lines used in this study include: *Tg(–.3drl:creERT2;cryaa:Venus)*, *Tg(– 3.2tbx1:creERT2;cryaa:Venus)*, *Tg(–1.5hsp70l:loxP-STOP-loxP-EGFP, cryaa:Venus)*, *Tg(crestin:creERT2;cryaa:YFP)*, *Tg(–3.5ubi:loxP-GFP-loxP-mCherry)*, and *Tg(– 4.9sox10:creERT2;actb2:loxP-BFP-loxP-DsRed)* (Felker et al., 2018; Giovannone et al., 2019; Kaufman et al., 2016; Mosimann et al., 2011; Mosimann et al., 2015). *Sox10:creERT2* fish were identified by PCR (Forward primer: 5’-TGCTGTTTCACTGGTTATGCGG-3’ and reverse primer: 5‘– TTGCCCCTGTTTCACTATCCAG-3’). Other transgenic fishes were screened by fluorescence markers–*cryaa:Venus, cryaa:YFP, ubi:Switch,* and *actb2:Switch*.

### Staging

Since experimental conditions, such as photoconversion, overnight shipment, and heat shock may affect the developmental speed of zebrafish embryos, embryonic stages were determined based on external morphology not on absolute time, with reference to the staging table provided by Kimmel et al., 1995.

### Photoconversion-based spatial lineage tracing

To prepare the DNA template for Kaede mRNA synthesis, the coding sequence of Kaede was subcloned from pME-Kaede vector into pCS2+ expression vector. Kaede mRNA was synthesized *in vitro* using mMESSAGE mMACHINE SP6 kit (AM1340, Invitrogen) following the manufacturer’s instructions. To obtain zebrafish embryos with ubiquitous Kaede expression, Kaede mRNA diluted at 100 ng/µl in RNase-free water containing 0.005% phenol red (P0290-100ML, Sigma) was injected into the cytoplasm of wild-type embryos at one-cell stage using an MPPI-3 microinjector (ASI) under a stereomicroscope (S9E, Leica). Injected embryos were incubated in the embryo medium (10% Hank’s with full strength calcium and magnesium: 13.7 mM NaCl, 0.54 mM KCl, 0.025 mM Na_2_HPO_4_, 0.044 mM KH_2_PO_4_, 1.3 mM CaCl_2_, 1.0 mM MgSO_4_, 0.42 mM NaHCO_3_). To photoconvert target embryonic regions at the 10–12 somite stage, we embedded embryos into 0.7% low gelling temperature agarose (50100, FMC BioProducts) in the embryo medium on a glass bottom dish (150680, Thermo Fisher Scientific) filled by phenol red-free L-15 medium (21083-027, Gibco). We then conducted photoconversion by illuminating 405 nm laser to the target regions for approximately 60 seconds on LSM510 Meta inverted confocal microscope (Zeiss). Labeled embryos were removed from the gel and incubated in six-well plates (657160, Greiner bio-one) filled with the embryo medium containing 0.003% 1-phenyl 2-thiourea (PTU; P7629, Sigma-Aldrich) at 28.5°C until 72 hpf. Labeled embryos were anesthetized with 0.17 mg/ml ethyl 3-aminobenzoate methanesulfonateMS-222 (E10521-50G, Sigma-Aldrich), embedded in 0.7% agarose gel, and imaged alive with the confocal microscope. All labeled results are listed in Supplementary file 1. For the consistency of the data, individuals with weak Kaede fluorescence or an excess amount of ectopic labeling (shown with asterisks in Supplementary file 1) were omitted from the analysis (individuals highlighted in gray in Supplementary file 1).

### Genetic lineage tracing

For heat-shock induction of *loxP* reporter cassettes driven by *hsp70l* promoter and activation of *CreERT2* in *drl:creERT2*– and *tbx1:creERT2*-lineages, we followed the previously reported procedures (Lalonde et al., 2022). For CreERT2 activation in the *crestin:creERT2*– and *sox10:creERT2*-lineage, we added 5 µM 4-OHT to the embryo medium containing the dechorionated embryos from shield stage to 24 hpf and 11 to 24 hpf, respectively. After the 4-OHT treatment, embryos were rinsed, raised to desired stages (72 or 96 hpf) in the embryo medium containing 0.003% PTU at 28.5°C, and fixed by 4% paraformaldehyde (PFA) in phosphate-buffered saline (PBS) at 4°C overnight.

### *In situ* hybridization

Probe sets for HCR RNA fluorescence *in situ* hybridization were designed to be complementary to target transcripts and synthesized by Molecular Instruments (Key Resources Table). For the whole-mount *in situ* HCR, fixed embryos were washed with PBST (PBS containing 0.1% Tween 20 [P20370-1.0, RPI Research Products International]) at least three times. Subsequent procedures followed the manufacturer’s instructions (MI-Protocol-RNAFISH-Zebrafish, Rev.#: 9) (Choi et al., 2018). After the HCR, nuclei and actin were stained in PBST containing 5 µg/ml 4′,6-diamidino-2– phenylindole dihydrochloride (DAPI; D9542, Sigma-Aldrich) and Alexa Fluor 488 phalloidin (diluted in 1/100; A12379, Life Technologies), respectively, without light exposure at room temperature for two overnights.

### Immunofluorescence

A duplexed *in situ* HCR and immunofluorescence has been performed as previously described (Ibarra-García-Padilla et al., 2021) with minor modifications. Briefly, we omitted the methanol and acetone permeabilization steps before the whole mount HCR from the original protocol. Embryos after *in situ* HCR were washed with PBST and soaked in blocking solution [5% heat inactivated (56°C, 30 min) sheep serum (S2263, Sigma-Aldrich) in PBST] at room temperature for 2 hours. The blocking solution was replaced by a primary antibody solution [anti-mCherry rabbit polyclonal antibody (GTX128508, GeneTex) diluted in 1/200 in blocking solution]. Then, we incubated embryos with the primary antibody solution at 4°C overnight. We washed the excess antibody solution by PBST and replaced the solution with a secondary antibody solution [anti–rabbit IgG Alexa Fluor 594 antibody (A11037, Invitrogen) diluted in 1/200]. Subsequently, we incubated embryos with a secondary antibody solution at 4°C overnight. The excess secondary antibody solution was removed and embryos were washed with PBST and mounted in the 0.7 % low-gelling temperature agarose gel in PBST. Finally, mounted samples were subjected to a graded series of glycerol/PBST and transferred into 75 % glycerol in PBST and imaged.

### Imaging

Bright-field and fluorescence whole-mount images were photographed an upright microscope MZ16 (Leica) equipped with a digital camera MC170HD (Leica). Confocal sections were acquired by LSM510 Meta and LSM 880 inverted confocal laser microscope equipped with a Plan-Apochromat 20x/0.8 M27 or C-Apochromat 40x/1.20W Korr UV-VIS-IR M27 objective (Zeiss). The whole-mount and confocal images were processed by ImageJ (NIH) and ZEN 3.5 software (Zeiss), respectively. 3D reconstructions from serial confocal images were performed by ZEN 3.5 software. Exported images in TIFF format were then assembled into figures in Adobe Photoshop (Adobe Systems). Three or more observers independently lead the conclusion, unless otherwise described.

### Anatomical term

Cleithrohyoid muscle: This muscle originates from the anteroventral potion of the cleithrum and inserts on the urohyal bone in the zebrafish (Diogo et al., 2008). This muscle is often referred to as “sternohyoideus” in the fish community (Winterbottom, 1973). Although there is little doubt that this muscle is homologous to the sternohyoid muscle of amniotes given its innervation (spinal nerves at the occipital level; Ma et al., 2010), embryonic origin (anterior somites; Minchin et al., 2013), and developmental environment (compare Figure 4 in this study with Adachi et al., 2020), the authors considered that the use of the term containing “sterno” for teleost fish that do not possess the sternum would make a confusion. Therefore, referring to the previously published synonym list of the fish sternohyoideus (Winterbottom, 1973), the authors use “cleithrohyoid muscle” that accurately reflects the anatomical connectivity of muscles and skeletons in zebrafish.

### Authors’ contributions

S. K. and T. N. conceived the project and designed the experiments. S. K. performed molecular, lineage tracing, and histological experiments. R. L. L., T. A. M., and C. M. conducted genetic lineage tracing experiments for the *drl:creERT2*, *tbx1:creERT2*, and *crestin:creERT2* lineages. S. K., R. L. L., C. M., and T. N. wrote the manuscript.

## Acknowledgements

This research was supported by the National Science Foundation under Grant IOS 2210072 and the Institutional support provided by the Rutgers University School of Arts and Sciences and the Human Genetics Institute of New Jersey (to T. N.). We thank Gage Crump and Claire Arata for providing *sox10:creERT2;actb2:Switch* embryos, Charles K. Kaufman for providing adult *crestin:creERT2* fish, Hannah Cohen and Kathleen Flaherty for maintaining zebrafish populations in the aquarium, and Kazutaka Hosoda for a helpful discussion.

## ORCIDs

Shunya Kuroda: 0000-0003-4820-2165

Robert L. Lalonde: 0000-0002-18030983

Christian Mosimann: 0000-0002-07492576

Tetsuya Nakamura: 0000-0003-0183-1685

## Data availability

Any imaging data generated in this study are available from the corresponding authors upon request.

## Additional files

Supplementary file 1: Data generated in photoconversion-based lineage tracing. Abbreviations are listed within the file.

## Figures and legends

**Figure 2—figure supplement 1.**
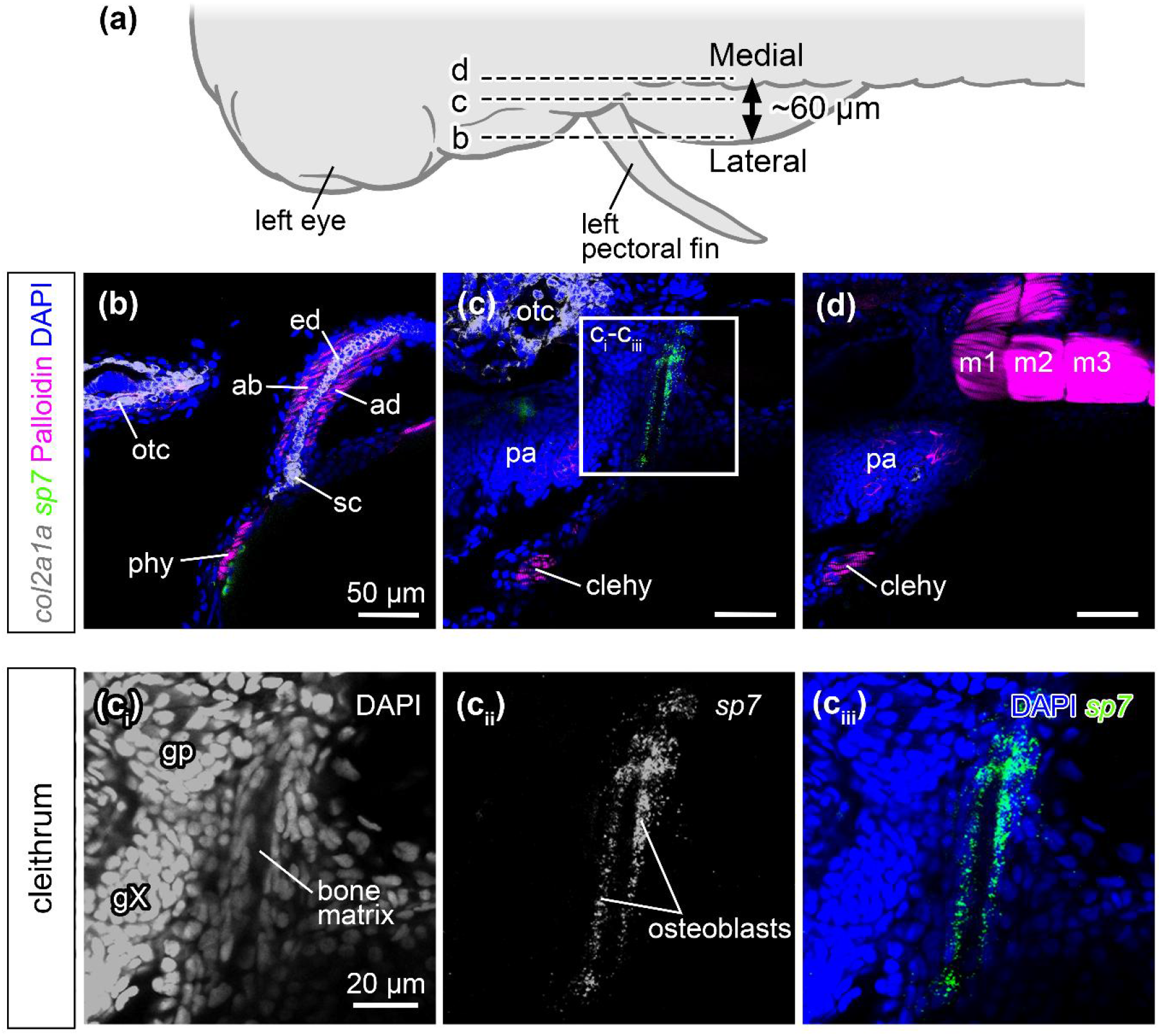
Histological and molecular reference of zebrafish pectoral region. **(a)** Dorsal view of the left half of zebrafish embryo at 72 hpf. Approximately 60 µm thickness regions in (a) were scanned along the lateral (b) to medial (d) direction by a confocal microscope. **(b–d)** Staining patterns of DAPI (nuclei), phalloidin (actin), *col2a1a* (cartilage), and *sp7*, (osteoblasts). The *cola2a1a* expressing cells (grey) exhibit a typical large, rounded, and tightly packed chondrocyte morphology in the endoskeletal disc and the scapulocoracoid (b). Skeletal muscles marked by phalloidin (magenta) show the typical histology of oriented striated muscle fibers in the pectoral fin adductor/abductor muscles, posterior hypaxial muscle, cleithrohyoid muscle, and myotomes (b, c). **(ci-ciii)** The enlarged view of the inset in (c) shows *sp7* positive cells (green) delineating the bone matrix of the cleithrum. ab, abductor muscle; ad, adductor muscle; clehy, cleithrohyoid muscle; ed, endoskeletal disc; gX, vagus ganglia; gp, posterior lateral line ganglion; m1–3, myotomes 1–3; otc, otic capsule; pa, pharyngeal arches; phy, posterior hypaxial muscle; sc, scapulocoracoid. Scale bars: (b–d) 50 µm, (ci–ciii) 20 µm.

**Figure 3—figure supplement 1.**
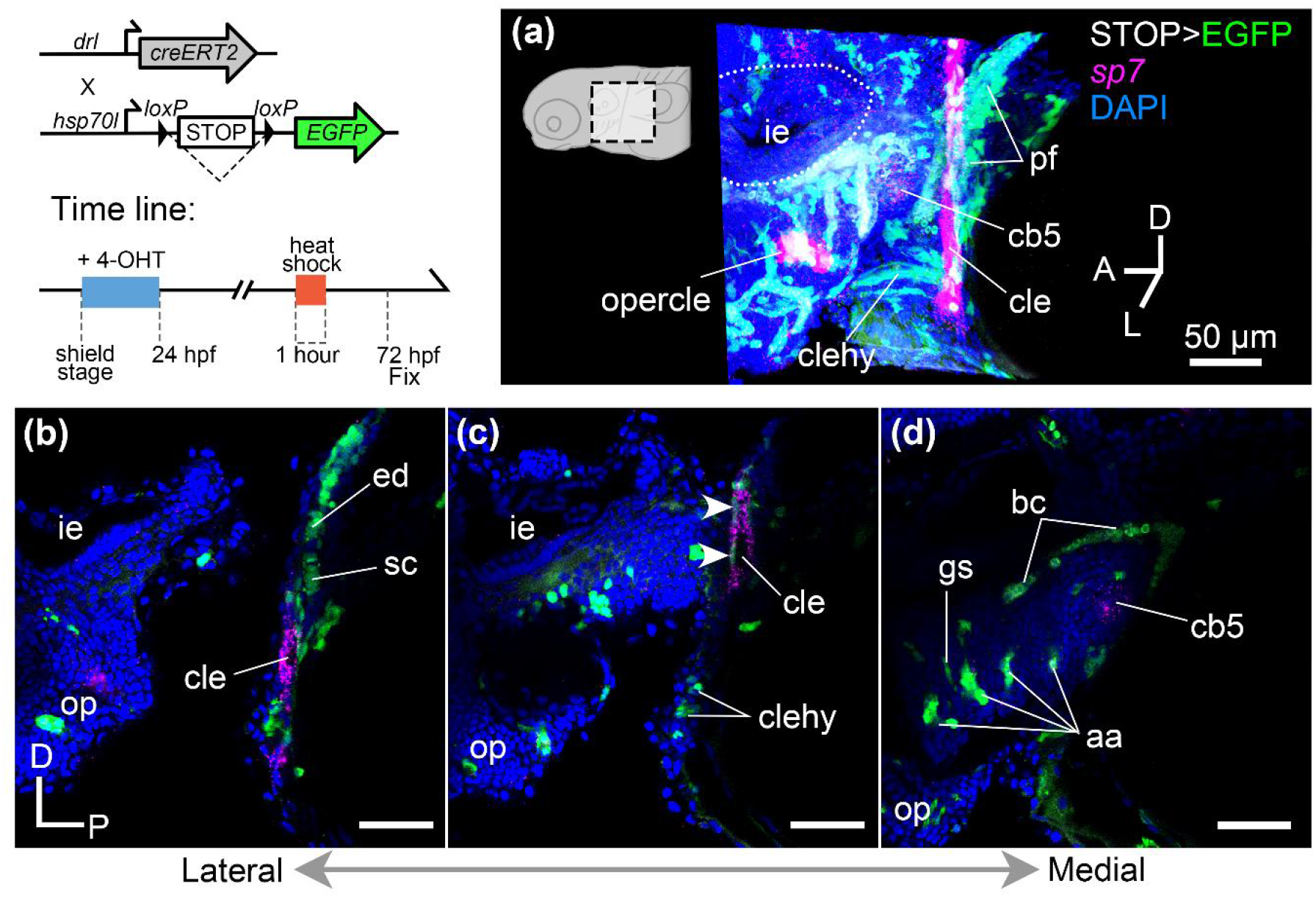
The *drl:creERT2* lineage contribution to the pectoral region. **(a)** Volumetric rendering of serial confocal images shows that *drl:creERT2* lineage with EGFP fluorescence (green) differentiates to *sp7*-positive osteoblasts (magenta) in the pectoral region of 72 hpf embryo. **(b–d)** Parasagittal confocal sections obtained at mediolaterally different levels [lateral (b) to medial (d)] show that *drl:creERT2* lineage cells are in the endoskeletal disc, scapulocoracoid, cleithrohyoid muscle, *sp7*-positive osteoblasts in the cleithrum (arrowheads), aortic arches, and endodermal epithelium in the gill slits. Green fluorescence in blood cells is likely autofluorescence. aa, aortic arches; bc, blood cells; cb5, ceratobranchial5; cle, cleithrum; clehy, cleithrohyoid; ed, endoskeletal disc; gs; gill slits; ie, inner ear; op, operculum; pf, pectoral fin; sc, scapulocoracoid. Scale bars: (a–d) 50 µm.

**Figure 4—figure supplement 1.**
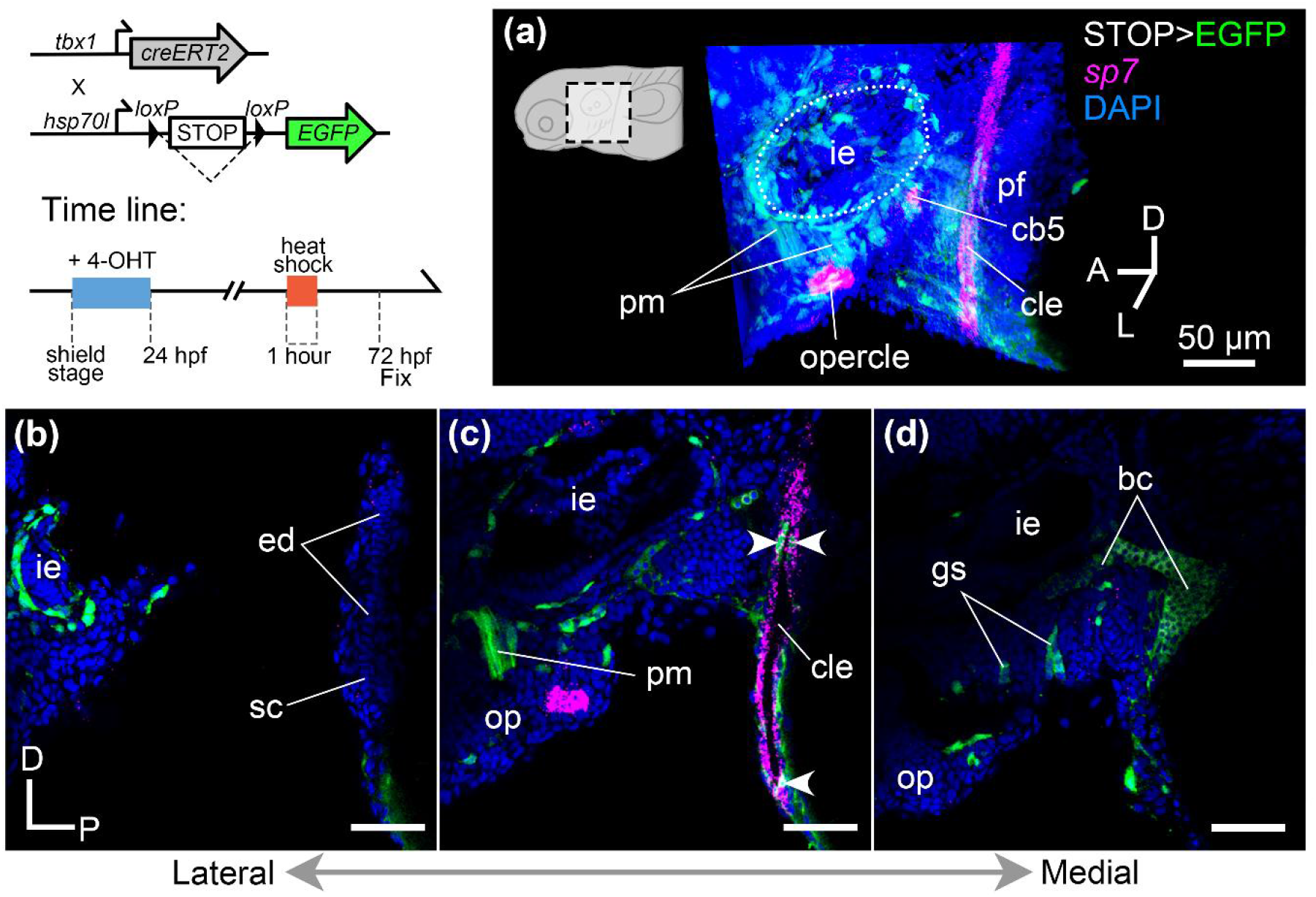
The *tbx1:creERT2* lineage contribution to the pectoral region. **(a)** Volumetric rendering of serial confocal images showed *tbx1:creERT2* lineage (EGFP, green) give rise to *sp7*-positive cells (magenta) in the pectoral region of 72 hpf embryo. **(b–d)** Parasagittal confocal sections at mediolaterally different levels [lateral (b) to medial (d)] show that labeled cells are in the mesenchymal cells surrounding the inner ear, pharyngeal musculature, *sp7*-positive osteoblasts in the cleithrum (arrowheads), and endodermal epithelium in the gill slits. Green fluorescence in blood cells is autofluorescence. bc, blood cells; cb5, ceratobranchial5; cle, cleithrum; ed, endoskeletal disc; gs; gill slits; ie, inner ear; op, operculum; pf, pectoral fin; pm, pharyngeal musculature; sc, scapulocoracoid. Scale bars: (a–d) 50 µm.

**Figure 5—figure supplement 1.**
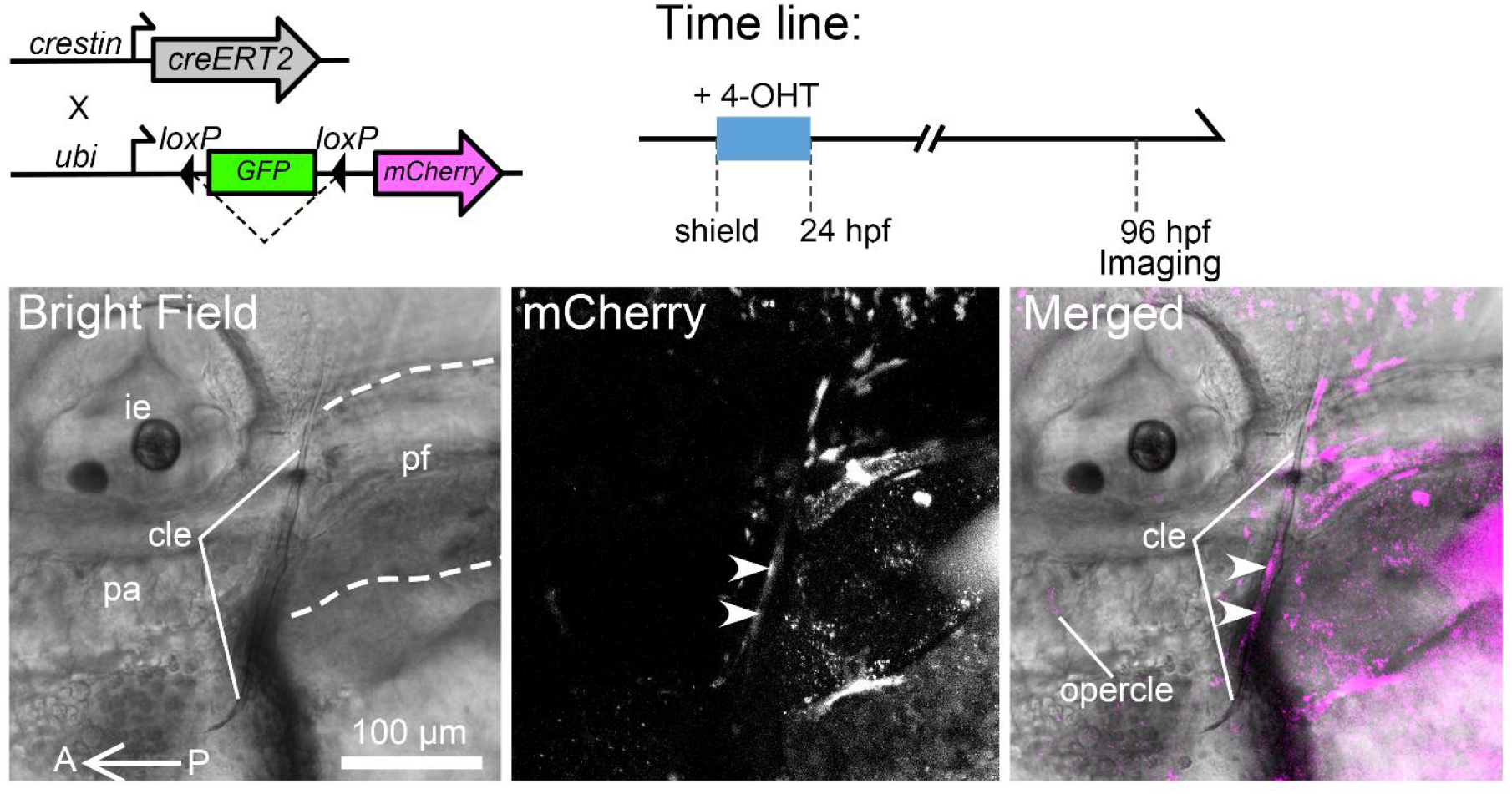
The *crestin:creERT2* lineage contribution to the cleithrum. Left lateral view of the *crestin:creERT2;ubi:Switch* embryo at 96 hpf. An overlayed image (right) of brightfield (left) and mCherry fluorescence projected along z-stacks (middle) shows that the *crestin:creERT2* lineage contributes to the cleithrum (arrowheads) and opercle. A white dashed line in the bright field image traces the outline of the pectoral fin. A, anterior; cle, cleithrum; ie, inner ear; op, operculum; P, posterior; pa, pharyngeal arches; pf, pectoral fin. Scale bar: 100 µm.

**Figure 5—figure supplement 2.**
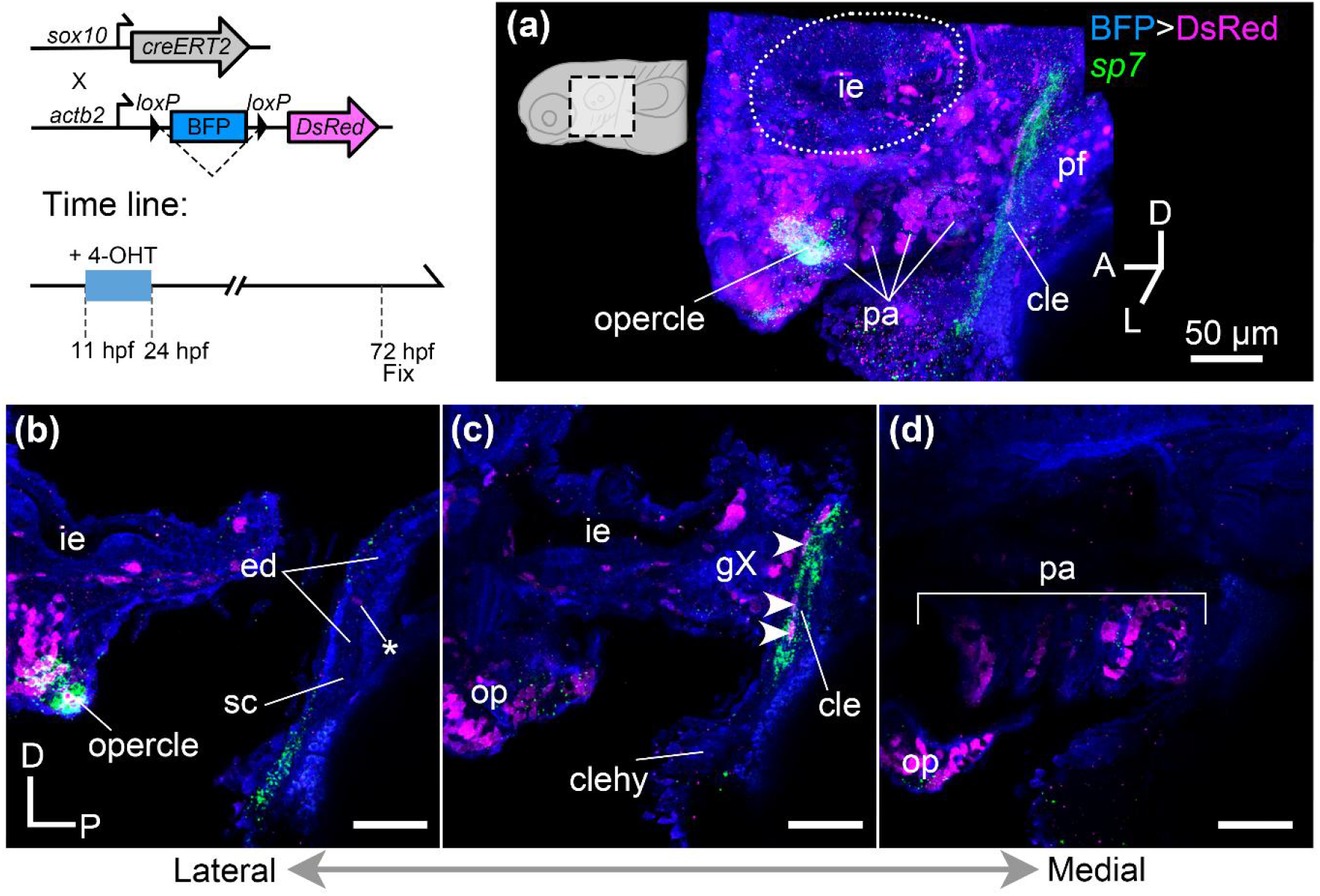
The *sox10:creERT2* lineage contribution to the pectoral region. **(a)** volumetric rendering of serial confocal images demonstrates that *sox10:creERT2* lineage marked by DsRed (magenta, immunofluorescence) generates *sp7*-positive osteoblasts (green, *in situ* HCR) in the pectoral region at 72 hpf. **(b–d)** Lateral confocal sections at mediolaterally different levels [lateral (b) to medial (d)] show that labeled cells are in the inner ear epithelium, operculum, pharyngeal mesenchyme, and *sp7* expressing osteoblasts in the anterior half of the cleithrum (arrowheads), and opercle bone. An ectopically labeled cartilaginous cell in the endoskeletal disc is indicated by an asterisk in panel (b). cle, cleithrum; clehy, cleithrohyoid muscle; ed, endoskeletal disc; gX, vagus ganglia; ie, inner ear; op, operculum; pa, pharyngeal arches; pf, pectoral fin; sc, scapulocoracoid. Scale bars: (a–d) 50 µm.

**Figure 6—figure supplement 1.**
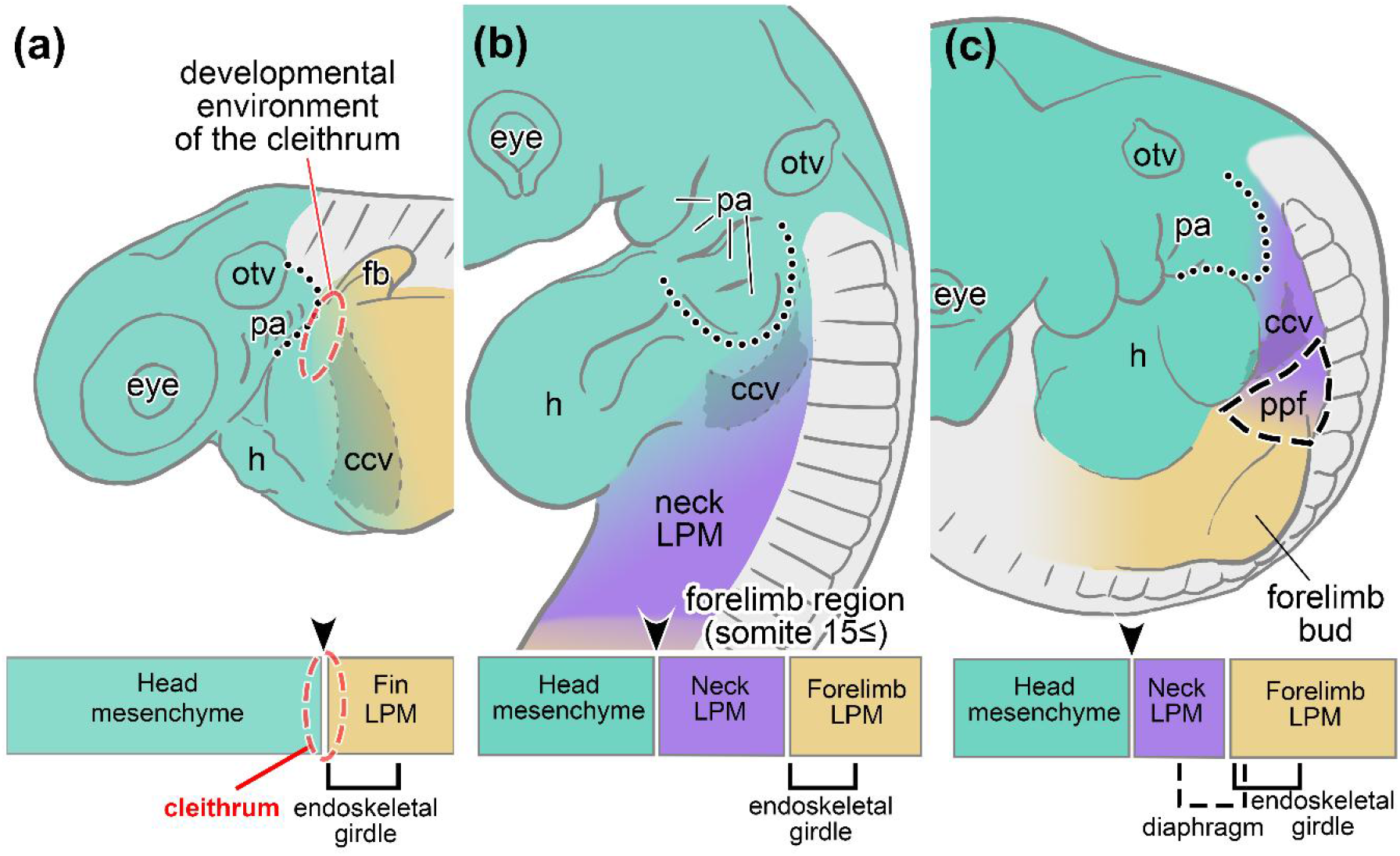
Comparison of the embryonic environment in the head and pectoral region between the animals with and without cleithrum. Schematic drawing of the left lateral view of **(a)** zebrafish, **(b)** chicken, **(c)** mouse embryos at 48 hpf, stage 18, and embryonic day 10–10.5, respectively. The posterior distribution limit of the head mesenchyme (combined cranial neural crest cells and CPM; colored in blue-green) is commonly located between the circumpharyngeal ridge (dotted lines) and common cardinal vein in gnathostomes. In contrast, the developmental environment of the cleithrum (encircled by red dashed line) is not identifiable in the long-necked amniotes such as chickens and mice due to the evolutionary insertion of the neck LPM (purple). In the mouse embryo, transitional region between the neck and forelimb LPM gives rise to the pleuroperitoneal fold (outlined by the dashed line). Arrowheads point the position of embryonic head/trunk interface. The distribution pattern of each embryonic cell population in the long-necked amniote is based on the following literature: chicken embryo (Lours-Calet et al., 2014; Nagashima et al., 2016), mouse embryo (Adachi et al., 2018; Heude et al., 2018; Sefton et al., 2018). Not to scale. ccv, common cardinal vein; fb, fin bud; h, heart; LPM, lateral plate mesoderm; otv, otic vesicle; pa, pharyngeal arches; ppf, pleuroperitoneal fold.

## References

1. Abrial, M., Paffett-Lugassy, N., Jeffrey, S., Jordan, D., O’Loughlin, E., Frederick, C. J., Burns, C. G., & Burns, C. E. 2017. TGF-β signaling is necessary and sufficient for pharyngeal arch artery angioblast formation. Cell Reports, 20(4), 973–983. https://doi.org/10.1016/j.celrep.2017.07.002

2. Adachi, N., Bilio, M., Baldini, A., & Kelly, R. G. 2020. Cardiopharyngeal mesoderm origins of musculoskeletal and connective tissues in the mammalian pharynx. Development, 147(3), dev185256. https://doi.org/10.1242/dev.185256

3. Adachi, N., Pascual-Anaya, J., Hirai, T., Higuchi, S., & Kuratani, S. 2018. Development of hypobranchial muscles with special reference to the evolution of the vertebrate neck. Zoological Letters, 4, 5. https://doi.org/10.1186/s40851-018-0087-x

4. Adachi, N., Robinson, M., Goolsbee, A., & Shubin, N. H. 2016. Regulatory evolution of *Tbx5* and the origin of paired appendages. Proceedings of the National Academy of Sciences, 113(36), 10115–10120. https://doi.org/10.1073/pnas.1609997113

5. Aoto, K., Sandell, L. L., Butler Tjaden, N. E., Yuen, K. C., Watt, K. E. N., Black, B. L., Durnin, M., & Trainor, P. A. 2015. *Mef2c-F10N* enhancer driven β-galactosidase (LacZ) and Cre recombinase mice facilitate analyses of gene function and lineage fate in neural crest cells. Developmental Biology, 402(1), 3–16. https://doi.org/10.1016/j.ydbio.2015.02.022

6. Arnold, J. S., Werling, U., Braunstein, E. M., Liao, J., Nowotschin, S., Edelmann, W., Hebert, J. M., & Morrow, B. E. 2006. Inactivation of *Tbx1* in the pharyngeal endoderm results in 22q11DS malformations. Development, 133(5), 977–987. https://doi.org/10.1242/dev.02264

7. Capellini, T. D., Vaccari, G., Ferretti, E., Fantini, S., He, M., Pellegrini, M., Quintana, L., Di Giacomo, G., Sharpe, J., Selleri, L., & Zappavigna, V. 2010. Scapula development is governed by genetic interactions of *Pbx1* with its family members and with *Emx2* via their cooperative control of *Alx1*. Development, 137(15), 2559–2569. https://doi.org/10.1242/dev.048819

8. Carney, T. J., Dutton, K. A., Greenhill, E., Delfino-Machín, M., Dufourcq, P., Blader, P., & Kelsh, R. N. 2006. A direct role for Sox10 in specification of neural crest-derived sensory neurons. Development, 133(23), 4619–4630. https://doi.org/10.1242/dev.02668

9. Choi, H. M. T., Schwarzkopf, M., Fornace, M. E., Acharya, A., Artavanis, G., Stegmaier, J., Cunha, A., & Pierce, N. A. 2018. Third-generation *in situ* hybridization chain reaction: multiplexed, quantitative, sensitive, versatile, robust. Development, 145(12), dev165753. https://doi.org/10.1242/dev.165753

10. Das, A., & Crump, J. G. 2012. Bmps and Id2a act upstream of Twist1 to restrict ectomesenchyme potential of the cranial neural crest. PLOS Genetics, 8(5), e1002710. https://doi.org/10.1371/journal.pgen.1002710

11. Diogo, R., Hinits, Y., & Hughes, S. M. 2008. Development of mandibular, hyoid and hypobranchial muscles in the zebrafish: homologies and evolution of these muscles within bony fishes and tetrapods. BMC Developmental Biology, 8(1), 24. https://doi.org/10.1186/1471-213X-8-24

12. Durland, J. L., Sferlazzo, M., Logan, M., & Burke, A. C. 2008. Visualizing the lateral somitic frontier in the Prx1Cre transgenic mouse. Journal of Anatomy, 212(5), 590–602. https://doi.org/10.1111/j.1469-7580.2008.00879.x

13. Dutton, J. R., Antonellis, A., Carney, T. J., Rodrigues, F. S. L. M., Pavan, W. J., Ward, A., & Kelsh, R. N. 2008. An evolutionarily conserved intronic region controls the spatiotemporal expression of the transcription factor Sox10. BMC Developmental Biology, 8(1), 105. https://doi.org/10.1186/1471-213X-8-105

14. Epperlein, H. H., Khattak, S., Knapp, D., Tanaka, E. M., & Malashichev, Y. B. 2012. Neural crest does not contribute to the neck and shoulder in the axolotl (*Ambystoma mexicanum*). PLoS One, 7(12), e52244. https://doi.org/10.1371/journal.pone.0052244

15. Etchevers, H. C., Dupin, E., & Le Douarin, N. M. 2019. The diverse neural crest: from embryology to human pathology. Development, 146(5). https://doi.org/10.1242/dev.169821

16. Evans, D. J., & Noden, D. M. 2006. Spatial relations between avian craniofacial neural crest and paraxial mesoderm cells. Developmental Dynamics, 235(5), 1310–1325. https://doi.org/10.1002/dvdy.20663

17. Fabian, P., & Crump, J. G. 2023. Reassessing the embryonic origin and potential of craniofacial ectomesenchyme. Seminars in Cell & Developmental Biology, 138, 45–53. https://doi.org/10.1016/j.semcdb.2022.03.018

18. Felker, A., Prummel, K. D., Merks, A. M., Mickoleit, M., Brombacher, E. C., Huisken, J., Panáková, D., & Mosimann, C. 2018. Continuous addition of progenitors forms the cardiac ventricle in zebrafish. Nature Communications, 9(1), 2001. https://doi.org/10.1038/s41467-018-04402-6

19. Gans, C., & Northcutt, R. G. 1983. Neural crest and the origin of vertebrates: a new head. Science, 220(4594), 268–273. https://doi.org/10.1126/science.220.4594.268

20. Gegenbaur, C. 1895. Clavicula und Cleithrum. Morphologisches Jahrbuch, 23, 1–20.

21. Gess, R., & Ahlberg, P. E. 2018. A tetrapod fauna from within the Devonian Antarctic Circle. Science, 360(6393), 1120–1124. https://doi.org/10.1126/science.aaq1645

22. Giovannone, D., Paul, S., Schindler, S., Arata, C., Farmer, D. J. T., Patel, P., Smeeton, J., & Crump, J. G. 2019. Programmed conversion of hypertrophic chondrocytes into osteoblasts and marrow adipocytes within zebrafish bones. Elife, 8, e42736. https://doi.org/10.7554/eLife.42736

23. Goodrich, E. S. 1930. Studies on the structure and development of vertebrates. London: McMillan.

24. Greil, A. 1913. Entwickelungsgeschichte des Kopfes und des Blutgefäßsystems von Ceratodus forsteri. II. Die epigenetischen Erwerbungen während der Stadien 39-48. Denkschriften der Medicinisch-Naturwissenschaftlichen Gesellschaft zu Jena, 4, 935–1492.

25. Hamilton, F. 1822. An account of the fishes found in the river Ganges and its branches. London: Edinburgh.

26. Hatta, K., Tsujii, H., & Omura, T. 2006. Cell tracking using a photoconvertible fluorescent protein. Nature Protocols, 1(2), 960–967. https://doi.org/10.1038/nprot.2006.96

27. Heude, E’., Shaikho, S., & Ekker, M. 2014. The *dlx5a/dlx6a* genes play essential roles in the early development of zebrafish median fin and pectoral structures. PLoS One, 9(5), e98505. https://doi.org/10.1371/journal.pone.0098505

28. Heude, E., Tesarova, M., Sefton, E. M., Jullian, E., Adachi, N., Grimaldi, A., Zikmund, T., Kaiser, J., Kardon, G., Kelly, R. G., & Tajbakhsh, S. 2018. Unique morphogenetic signatures define mammalian neck muscles and associated connective tissues. Elife, 7, e40179. https://doi.org/10.7554/eLife.40179

29. Higashiyama, H., Hirasawa, T., Oisi, Y., Sugahara, F., Hyodo, S., Kanai, Y., & Kuratani, S. 2016. On the vagal cardiac nerves, with special reference to the early evolution of the head–trunk interface. Journal of Morphology, 277(9), 1146–1158. https://doi.org/10.1002/jmor.20563

30. Hildebrand, D. G. C., Cicconet, M., Torres, R. M., Choi, W., Quan, T. M., Moon, J., Wetzel, A. W., Scott Champion, A., Graham, B. J., Randlett, O., Plummer, G. S., Portugues, R., Bianco, I. H., Saalfeld, S., Baden, A. D., Lillaney, K., Burns, R., Vogelstein, J. T., Schier, A. F., Lee, W.-C. A., Jeong, W.-K., Lichtman, J. W., & Engert, F. 2017. Whole-brain serial-section electron microscopy in larval zebrafish. Nature, 545(7654), 345–349. https://doi.org/10.1038/nature22356

31. Hirasawa, T., Cupello, C., Brito, P. M., Yabumoto, Y., Isogai, S., Hoshino, M., & Uesugi, K. 2021. Development of the pectoral lobed fin in the Australian lungfish *Neoceratodus forsteri*. Frontiers in Ecology and Evolution, 9(484). https://doi.org/10.3389/fevo.2021.679633

32. Hirasawa, T., Fujimoto, S., & Kuratani, S. 2016. Expansion of the neck reconstituted the shoulder-diaphragm in amniote evolution. Development, Growth & Differentiation, 58(1), 143–153. https://doi.org/10.1111/dgd.12243

33. Hirasawa, T., & Kuratani, S. 2013. A new scenario of the evolutionary derivation of the mammalian diaphragm from shoulder muscles. Journal of Anatomy, 222(5), 504–517. https://doi.org/10.1111/joa.12037

34. Howard IV, A. G. A., Baker, P. A., Ibarra-García-Padilla, R., Moore, J. A., Rivas, L. J., Tallman, J. J., Singleton, E. W., Westheimer, J. L., Corteguera, J. A., & Uribe, R. A. 2021. An atlas of neural crest lineages along the posterior developing zebrafish at single-cell resolution. Elife, 10, e60005. https://doi.org/10.7554/eLife.60005

35. Ibarra-García-Padilla, R., Howard, A. G. A., Singleton, E. W., & Uribe, R. A. 2021. A protocol for whole-mount immuno-coupled hybridization chain reaction (WICHCR) in zebrafish embryos and larvae. STAR Protocols, 2(3), 100709. https://doi.org/10.1016/j.xpro.2021.100709

36. Janvier, P. 1996. Early Vertebrates. Oxford: Oxford University Press.

37. Janvier, P., Percy, L. R., & Potter, I. C. 1991. The arrangement of the heart chambers and associated blood vessels in the Devonian osteostracan *Norselaspis glacialis*. A reinterpretation based on recent studies of the circulatory system in lampreys. Journal of Zoology, 223(4), 567–576. https://doi.org/10.1111/j.1469-7998.1991.tb04388.x

38. Jollie, M. 1962. Chordate Morphology. London: Chapman & Hall.

39. Kague, E., Gallagher, M., Burke, S., Parsons, M., Franz-Odendaal, T., & Fisher, S. 2012. Skeletogenic fate of zebrafish cranial and trunk neural crest. PLoS One, 7(11), e47394. https://doi.org/10.1371/journal.pone.0047394

40. Kaufman, C. K., Mosimann, C., Fan, Z. P., Yang, S., Thomas, A. J., Ablain, J., Tan, J. L., Fogley, R. D., van Rooijen, E., Hagedorn, E. J., Ciarlo, C., White, R. M., Matos, D. A., Puller, A.-C., Santoriello, C., Liao, E. C., Young, R. A., & Zon, L. I. 2016. A zebrafish melanoma model reveals emergence of neural crest identity during melanoma initiation. Science, 351(6272), aad2197. https://doi.org/10.1126/science.aad2197

42. Kimmel, C. B., Ballard, W. W., Kimmel, S. R., Ullmann, B., & Schilling, T. F. 1995. Stages of embryonic development of the zebrafish. Developmental Dynamics, 203(3), 253–310. https://doi.org/10.1002/aja.1002030302

43. Kingsley, J. S. 1917. Outlines of comparative antomy of vertebrates (second ed.). Philadelphia: P. Blakston’s son & Co.

44. Kuratani, S. 1997. Spatial distribution of postotic crest cells defines the head/trunk interface of the vertebrate body: embryological interpretation of peripheral nerve morphology and evolution of the vertebrate head. Anatomy and Embryology, 195(1), 1–13. https://doi.org/10.1007/s004290050020

45. Kuratani, S., & Ahlberg, P. E. 2018. Evolution of the vertebrate neurocranium: problems of the premandibular domain and the origin of the trabecula. Zoological Letters, 4(1), 1. https://doi.org/10.1186/s40851-017-0083-6

46. Lalonde, R. L., Kemmler, C. L., Riemslagh, F. W., Aman, A. J., Kresoja-Rakic, J., Moran, H. R., Nieuwenhuize, S., Parichy, D. M., Burger, A., & Mosimann, C. 2022. Heterogeneity and genomic loci of ubiquitous transgenic Cre reporter lines in zebrafish. Developmental Dynamics, 251(10), 1754–1773. https://doi.org/10.1002/dvdy.499

47. Lee, R. T. H., Knapik, E. W., Thiery, J. P., & Carney, T. J. 2013. An exclusively mesodermal origin of fin mesenchyme demonstrates that zebrafish trunk neural crest does not generate ectomesenchyme. Development, 140(14), 2923–2932. https://doi.org/10.1242/dev.093534

48. Lescroart, F., Dumas, C. E., Adachi, N., & Kelly, R. G. 2022. Emergence of heart and branchiomeric muscles in cardiopharyngeal mesoderm. Experimental Cell Research, 410(1), 112931. https://doi.org/10.1016/j.yexcr.2021.112931

49. Lours-Calet, C., Alvares, L. E., El-Hanfy, A. S., Gandesha, S., Walters, E. H., Sobreira, D. R., Wotton, K. R., Jorge, E. C., Lawson, J. A., Kelsey Lewis, A., Tada, M., Sharpe, C., Kardon, G., & Dietrich, S. 2014. Evolutionarily conserved morphogenetic movements at the vertebrate head-trunk interface coordinate the transport and assembly of hypopharyngeal structures. Developmental Biology, 390(2), 231–246. https://doi.org/10.1016/j.ydbio.2014.03.003

50. Lours, C., & Dietrich, S. 2005. The dissociation of the Fgf-feedback loop controls the limbless state of the neck. Development, 132(24), 5553–5564. https://doi.org/10.1242/dev.02164

51. Lyson, T. R., Bhullar, B.-A. S., Bever, G. S., Joyce, W. G., de Queiroz, K., Abzhanov, A., & Gauthier, J. A. 2013. Homology of the enigmatic nuchal bone reveals novel reorganization of the shoulder girdle in the evolution of the turtle shell. Evolution & Development, 15(5), 317–325. https://doi.org/10.1111/ede.12041

52. Müller, J., Scheyer, T. M., Head, J. J., Barrett, P. M., Werneburg, I., Ericson, P. G. P., Pol, D., & Sánchez-Villagra, M. R. 2010. Homeotic effects, somitogenesis and the evolution of vertebral numbers in recent and fossil amniotes. Proceedings of the National Academy of Sciences, 107(5), 2118–2123. https://doi.org/10.1073/pnas.0912622107

53. Ma, L.-H., Gilland, E., Bass, A. H., & Baker, R. 2010. Ancestry of motor innervation to pectoral fin and forelimb. Nature Communications, 1(1), 49. https://doi.org/10.1038/ncomms1045

54. MacDougall, M. J., Modesto, S. P., & Botha-Brink, J. 2013. The postcranial skeleton of the Early Triassic parareptile *Sauropareion anoplus*, with a discussion of possible life history. Acta Palaeontologica Polonica, 58(4), 737–749. https://doi.org/10.4202/app.2011.0099

55. Mao, Q., Stinnett, H. K., & Ho, R. K. 2015. Asymmetric cell convergence-driven zebrafish fin bud initiation and pre-pattern requires Tbx5a control of a mesenchymal Fgf signal. Development, 142(24), 4329–4339. https://doi.org/10.1242/dev.124750

56. Masselink, W., Cole, N. J., Fenyes, F., Berger, S., Sonntag, C., Wood, A., Nguyen, P. D., Cohen, N., Knopf, F., Weidinger, G., Hall, T. E., & Currie, P. D. 2016. A somitic contribution to the apical ectodermal ridge is essential for fin formation. Nature, 535(7613), 542–546. https://doi.org/10.1038/nature18953

57. Matsuoka, T., Ahlberg, P. E., Kessaris, N., Iannarelli, P., Dennehy, U., Richardson, W. D., McMahon, A. P., & Koentges, G. 2005. Neural crest origins of the neck and shoulder. Nature, 436(7049), 347–355. https://doi.org/10.1038/nature03837

58. McCarroll, M. N., Lewis, Z. R., Culbertson, M. D., Martin, B. L., Kimelman, D., & Nechiporuk, A. V. 2012. Graded levels of Pax2a and Pax8 regulate cell differentiation during sensory placode formation. Development, 139(15), 2740–2750. https://doi.org/10.1242/dev.076075

59. McGonnell, I. M. 2001. The evolution of the pectoral girdle. Journal of Anatomy, 199(1-2), 189–194. https://doi.org/10.1046/j.1469-7580.2001.19910189.x

60. McGonnell, I. M., McKay, I. J., & Graham, A. 2001. A population of caudally migrating cranial neural crest cells: Functional and evolutionary implications. Developmental Biology, 236(2), 354–363. https://doi.org/10.1006/dbio.2001.0330

61. Mercader, N. 2007. Early steps of paired fin development in zebrafish compared with tetrapod limb development. *Development*, Growth & Differentiation, 49(6), 421–437. https://doi.org/10.1111/j.1440-169X.2007.00942.x

62. Minchin, J. E. N., Williams, V. C., Hinits, Y., Low, S., Tandon, P., Fan, C.-M., Rawls, J. F., & Hughes, S. M. 2013. Oesophageal and sternohyal muscle fibres are novel Pax3-dependent migratory somite derivatives essential for ingestion. Development, 140(14), 2972–2984. https://doi.org/10.1242/dev.090050

63. Mongera, A., Singh, A. P., Levesque, M. P., Chen, Y.-Y., Konstantinidis, P., & Nüsslein-Volhard, C. 2013. Genetic lineage labeling in zebrafish uncovers novel neural crest contributions to the head, including gill pillar cells. Development, 140(4), 916–925. https://doi.org/10.1242/dev.091066

64. Montero-Balaguer, M., Lang, M. R., Sachdev, S. W., Knappmeyer, C., Stewart, R. A., De La Guardia, A., Hatzopoulos, A. K., & Knapik, E. W. 2006. The mother superior mutation ablates *foxd3* activity in neural crest progenitor cells and depletes neural crest derivatives in zebrafish. Developmental Dynamics, 235(12), 3199–3212. https://doi.org/10.1002/dvdy.20959

65. Mosimann, C., Kaufman, C. K., Li, P., Pugach, E. K., Tamplin, O. J., & Zon, L. I. 2011. Ubiquitous transgene expression and Cre-based recombination driven by the *ubiquitin* promoter in zebrafish. Development, 138(1), 169–177. https://doi.org/10.1242/dev.059345

66. Mosimann, C., Panáková, D., Werdich, A. A., Musso, G., Burger, A., Lawson, K. L., Carr, L. A., Nevis, K. R., Sabeh, M. K., Zhou, Y., Davidson, A. J., DiBiase, A., Burns, C. E., Burns, C. G., MacRae, C. A., & Zon, L. I. 2015. Chamber identity programs drive early functional partitioning of the heart. Nature Communications, 6(1), 8146. http://doi.org/10.1038/ncomms9146

67. Nagashima, H., Sugahara, F., Watanabe, K., Shibata, M., Chiba, A., & Sato, N. 2016. Developmental origin of the clavicle, and its implications for the evolution of the neck and the paired appendages in vertebrates. Journal of Anatomy, 229(4), 536–548. https://doi.org/10.1111/joa.12502

68. Naumann, B., Warth, P., Olsson, L., & Konstantinidis, P. 2017. The development of the cucullaris muscle and the branchial musculature in the Longnose Gar, (*Lepisosteus osseus*, Lepisosteiformes, Actinopterygii) and its implications for the evolution and development of the head/trunk interface in vertebrates. Evolution & Development, 19(6), 263–276. https://doi.org/10.1111/ede.12239

69. Nomaru, H., Liu, Y., De Bono, C., Righelli, D., Cirino, A., Wang, W., Song, H., Racedo, S. E., Dantas, A. G., Zhang, L., Cai, C.-L., Angelini, C., Christiaen, L., Kelly, R. G., Baldini, A., Zheng, D., & Morrow, B. E. 2021. Single cell multi-omic analysis identifies a *Tbx1*-dependent multilineage primed population in murine cardiopharyngeal mesoderm. Nature Communications, 12(1), 6645. https://doi.org/10.1038/s41467-021-26966-6

70. Onimaru, K., Shoguchi, E., Kuratani, S., & Tanaka, M. 2011. Development and evolution of the lateral plate mesoderm: Comparative analysis of amphioxus and lamprey with implications for the acquisition of paired fins. Developmental Biology, 359(1), 124–136. https://doi.org/10.1016/j.ydbio.2011.08.003

71. Piekarski, N., & Olsson, L. 2011. A somitic contribution to the pectoral girdle in the axolotl revealed by long-term fate mapping. Evolution & Development, 13(1), 47–57. https://doi.org/10.1111/j.1525-142X.2010.00455.x

72. Prummel, K. D., Crowell, H. L., Nieuwenhuize, S., Brombacher, E. C., Daetwyler, S., Soneson, C., Kresoja-Rakic, J., Kocere, A., Ronner, M., Ernst, A., Labbaf, Z., Clouthier, D. E., Firulli, A. B., Sánchez-Iranzo, H., Naganathan, S. R., O’Rourke, R., Raz, E., Mercader, N., Burger, A., Felley-Bosco, E., Huisken, J., Robinson, M. D., & Mosimann, C. 2022. Hand2 delineates mesothelium progenitors and is reactivated in mesothelioma. Nature Communications, 13(1), 1677. https://doi.org/10.1038/s41467-022-29311-7

73. Prummel, K. D., Nieuwenhuize, S., & Mosimann, C. 2020. The lateral plate mesoderm. Development, 147(12). https://doi.org/10.1242/dev.175059

74. Romer, A. S. 1997. Osteology of the reptiles (Rprint ed.). Florida: Krieger Publishing Company.

75. Romer, A. S., & Parsons, T. S. 1986. The vertebrate body (6th ed.). Philadelphia: Saunders College.

76. Sagarin, K. A., Redgrave, A. C., Mosimann, C., Burke, A. C., & Devoto, S. H. 2019. Anterior trunk muscle shows mix of axial and appendicular developmental patterns. Developmental Dynamics, 248(10), 961–968. https://doi.org/10.1002/dvdy.95

77. Sato, M., & Yost, H. J. 2003. Cardiac neural crest contributes to cardiomyogenesis in zebrafish. Developmental Biology, 257(1), 127–139. https://doi.org/10.1016/S0012-1606(03)00037-X

78. Schneider, R. A. 1999. Neural crest can form cartilages normally derived from mesoderm during development of the avian head skeleton. Developmental Biology, 208(2), 441–455. https://doi.org/10.1006/dbio.1999.9213

79. Sefton, E. M., Gallardo, M., & Kardon, G. 2018. Developmental origin and morphogenesis of the diaphragm, an essential mammalian muscle. Developmental Biology, 440(2), 64–73. https://doi.org/10.1016/j.ydbio.2018.04.010

80. Sefton, E. M., Gallardo, M., Tobin, C. E., Collins, B. C., Colasanto, M. P., Merrell, A. J., & Kardon, G. 2022. Fibroblast-derived *Hgf* controls recruitment and expansion of muscle during morphogenesis of the mammalian diaphragm. Elife, 11, e74592. https://doi.org/10.7554/eLife.74592

81. Sefton, E. M., Piekarski, N., & Hanken, J. 2015. Dual embryonic origin and patterning of the pharyngeal skeleton in the axolotl (*Ambystoma mexicanum*). Evolution & Development, 17(3), 175–184. https://doi.org/10.1111/ede.12124

82. Svestak, M. S., Božičević, V., Bakarić, R., Dunjko, V., & Domazet-Lošo, T. 2013. Phylostratigraphic profiles reveal a deep evolutionary history of the vertebrate head sensory systems. Frontiers in Zoology, 10(1), 18. https://doi.org/10.1186/1742-9994-10-18

83. Shearman, R. M., Tulenko, F. J., & Burke, A. C. 2011. 3D reconstructions of quail-chick chimeras provide a new fate map of the avian scapula. Developmental Biology, 355(1), 1–11. https://doi.org/10.1016/j.ydbio.2011.03.032

84. Sleight, V. A., & Gillis, J. A. 2020. Embryonic origin and serial homology of gill arches and paired fins in the skate, *Leucoraja erinacea*. Elife, 9, e60635. https://doi.org/10.7554/eLife.60635

85. Stundl, J., Pospisilova, A., Matějková, T., Psenicka, M., Bronner, M. E., & Cerny, R. 2020. Migratory patterns and evolutionary plasticity of cranial neural crest cells in ray-finned fishes. Developmental Biology, 467(1), 14–29. https://doi.org/10.1016/j.ydbio.2020.08.007

86. Tabler, J. M., Rigney, M. M., Berman, G. J., Gopalakrishnan, S., Heude, E., Al-lami, H. A., Yannakoudakis, B. Z., Fitch, R. D., Carter, C., Vokes, S., Liu, K. J., Tajbakhsh, S., Egnor, S. E. R., & Wallingford, J. B. 2017. Cilia-mediated Hedgehog signaling controls form and function in the mammalian larynx. Elife, 6, e19153. https://doi.org/10.7554/eLife.19153

87. Talbot, J. C., Teets, E. M., Ratnayake, D., Duy, P. Q., Currie, P. D., & Amacher, S. L. 2019. Muscle precursor cell movements in zebrafish are dynamic and require Six family genes. Development, 146(10). https://doi.org/10.1242/dev.171421

88. Teng, C. S., Cavin, L., Maxson, R. E. J., Sánchez-Villagra, M. R., & Crump, J. G. 2019. Resolving homology in the face of shifting germ layer origins: Lessons from a major skull vault boundary. Elife, 8, e52814. https://doi.org/10.7554/eLife.52814

89. Trinajstic, K., Long, J. A., Sanchez, S., Boisvert, C. A., Snitting, D., Tafforeau, P., Dupret, V., Clement, A. M., Currie, P. D., Roelofs, B., Bevitt, J. J., Lee, M. S. Y., & Ahlberg, P. E. 2022. Exceptional preservation of organs in Devonian placoderms from the Gogo lagerstätte. Science, 377(6612), 1311–1314. https://doi.org/10.1126/science.abf3289

90. Trinajstic, K., Sanchez, S., Dupret, V., Tafforeau, P., Long, J., Young, G., Senden, T., Boisvert, C., Power, N., & Ahlberg, P. E. 2013. Fossil musculature of the most primitive jawed vertebrates. Science, 341(6142), 160–164. https://doi.org/10.1126/science.1237275

91. Valasek, P., Theis, S., Krejci, E., Grim, M., Maina, F., Shwartz, Y., Otto, A., Huang, R., & Patel, K. 2010. Somitic origin of the medial border of the mammalian scapula and its homology to the avian scapula blade. Journal of Anatomy, 216(4), 482–488. https://doi.org/10.1111/j.1469-7580.2009.01200.x

92. Walker, C. 1999. Haploid screens and gamma-ray mutagenesis. Methods Cell Biol, 60, 43–70. 10.1016/s0091-679x(08)61893-2

93. Wang, W., Niu, X., Stuart, T., Jullian, E., Mauck, W. M., Kelly, R. G., Satija, R., & Christiaen, L. 2019. A single-cell transcriptional roadmap for cardiopharyngeal fate diversification. Nature Cell Biology, 21(6), 674–686. https://doi.org/10.1038/s41556-019-0336-z

94. Williston, S. W. 1925. The osteology of the reptiles (W. K. Gregory Ed.). Cambridge: Harvard University Press.

95. Winterbottom, R. 1973. A descriptive synonymy of the striated muscles of the Teleostei. Proceedings of the Academy of Natural Sciences of Philadelphia, 125, 225–317. Retrieved from http://www.jstor.org/stable/4064691

